# Regulation of Human PINK1 ubiquitin kinase by Serine167, Serine228 and Cysteine412 phosphorylation

**DOI:** 10.1101/2023.03.31.534916

**Authors:** Andrew D. Waddell, Hina Ojha, Shalini Agarwal, Christopher J. Clarke, Ana Terriente-Felix, Houjiang Zhou, Poonam Kakade, Axel Knebel, Andrew M. Shaw, Robert Gourlay, Joby Varghese, Renata F. Soares, Rachel Toth, Thomas Macartney, Patrick A. Eyers, Nick Morrice, Richard Bayliss, Alexander J. Whitworth, Claire E. Eyers, Miratul M. K. Muqit

## Abstract

Loss-of-function mutations in the human PINK1 kinase (*h*PINK1) are causative of early-onset Parkinson’s disease (PD). Activation of *h*PINK1 induces phosphorylated ubiquitin to initiate removal of damaged mitochondria by autophagy. Previously we solved the structure of the insect PINK1 orthologue, *Tribolium castaneum* PINK1, and showed that autophosphorylation of Ser205 was critical for ubiquitin interaction and phosphorylation (Kumar, Tamjar, Waddell et al., 2017). Here we report new findings on the regulation of *h*PINK1 by phosphorylation. We reconstitute *E. coli* expressed *h*PINK1 activity *in vitro* by direct incorporation of phosphoserine at the equivalent site Serine 228 (pSer228), providing direct evidence for a role for Ser228 phosphorylation in *h*PINK1 activation. Furthermore, using mass spectrometry, we identify six novel Ser/Thr autophosphorylation sites including regulatory Serine167 phosphorylation (pSer167), which in addition to pSer228 is required for ubiquitin recognition and phosphorylation. Strikingly, we also detect phosphorylation of a conserved Cysteine412 (pCys412) residue in the *h*PINK1 activation segment. Structural modelling suggests that pCys412 inhibits ubiquitin recognition and we demonstrate that mutation of Cys412 to Ala renders *h*PINK1 more active towards ubiquitin when expressed in human cells. These results outline new insights into *h*PINK1 activation by pSer167 and pSer228 and a novel inhibitory mechanism mediated by pCys412. These findings will aid in the development of small molecule activators of *h*PINK1.

## Introduction

Autosomal recessive mutations in PTEN-induced kinase 1 (PINK1) are the second most frequent cause of early-onset Parkinson’s disease (PD) [1]. PINK1 functions as a master regulator of mitochondrial quality control, promoting ubiquitin-dependent elimination of damaged mitochondria *via* autophagy (mitophagy) [2–4]. As the only known mitochondrial-localised protein kinase, PINK1 is constitutively targeted to mitochondria through an N-terminal import sequence, where it is rapidly turned over following consecutive processing by matrix and PARL proteases and therefore inactive under basal conditions [5–10]. However, upon mitochondrial depolarization of the inner membrane that can be induced artificially by mitochondrial uncoupling agents (*e.g.* carbonyl cyanide *m*-chlorophenyl hydrazone (CCCP)), PINK1 is stabilised and activated at the Translocase of Outer Membrane (TOM) complex where it phosphorylates its substrates Parkin and ubiquitin at an equivalent Serine65 (Ser65) residue [11–16]. This results in activation of Parkin via a feed-forward mechanism triggering mitophagy [17–19].

The molecular mechanism of human PINK1 (*h*PINK1) activation remains to be elucidated and has been hampered by low *in vitro* activity of recombinant *h*PINK1 expressed in *E. coli* or insect cells or immunoprecipitated from human cells [13, 20]. Strikingly, recombinant PINK1 protein from two insect orthologues *Tribolium castaneum* (*Tc*PINK1) and *Pediculus humanus corporis* (*Phc*PINK1) exhibit robust catalytic activity *in vitro* and autophosphorylate at a key Serine residue equivalent to human Serine228 (Ser228) [20]. Structural and biochemical analysis of *Tc*PINK1 and *Phc*PINK1 has experimentally defined Ser228 phosphorylation (pSer228) as essential for PINK1 substrate recognition via stabilisation of a unique loop insertion (Insertion 3; Ins3) within its catalytic domain [21–23]. Two groups recently solved structures of active insect PINK1 in a symmetrical dimer mediated by another loop insertion (Insertion 2; Ins2), resulting in Ser228 being positioned at the active site of the other kinase thereby enabling trans-autophosphorylation of Ser228 [24, 25]. In human cells, we recently found that *h*PINK1 recruitment to the TOM complex is a pre-requisite for Ser228 autophosphorylation using a specific anti-phospho-Ser228 PINK1 antibody [26].

Herein we perform reconstitution experiments of recombinant *h*PINK1 expressed in *E. coli* combined with gene codon expansion to demonstrate that pSer228 is sufficient to activate *h*PINK1 and elaborate the first catalytic assay of *h*PINK1 *in vitro*. We further undertake mutagenesis analysis of Ser228 in cell-based studies to demonstrate a critical role for pSer228 in *h*PINK1 activation following mitochondrial depolarisation. Employing targeted mass spectrometry (MS)-based phosphosite mapping of human *h*PINK1 in cells we identified six novel Ser/Thr autophosphorylation sites (pSer123, pSer161, pSer167, pThr185, pSer187, pSer245) in addition to previously reported pSer228, pThr257 and pSer284 sites. We demonstrate that pSer167 is required for ubiquitin substrate recognition and phosphorylation representing a new regulatory site in addition to pSer228. Strikingly, we also unambiguously detect phosphorylation of a conserved cysteine residue in the *h*PINK1 activation segment, Cysteine412 (pCys412). Structural modelling suggests that pCys412 inhibits ubiquitin recognition and we demonstrate that mutation of Cys412 to Ala renders *h*PINK1 more active towards ubiquitin when expressed in human cells. This systematic analysis of *h*PINK1 phosphorylation sites provides new fundamental insights into the regulation of *h*PINK1 and will aid in the development of small molecule activators of *h*PINK1.

## Results & Discussion

### Role of Ser228 autophosphorylation for *h*PINK1 activation *in vitro* and in cells

Ser228 is highly conserved across species, from mammals through to invertebrates (Figure 1a). Autophosphorylation of this residue is a known event during the activation of insect PINK1 [20], with crystal structures and HDX-MS showing that pSer228 induces a conformational change in Ins3, which is a major substrate-recognition element for ubiquitin or the ubiquitin-like domain (Ubl) of Parkin (Figure 1b) [21–23]. Mutations of insect PINK1 Ser228 to Ala or deletions or mutations of Ins3 therefore completely prevent ubiquitin and Parkin phosphorylation *in vitro* [21–23]. We generated an AlphaFold model of *h*PINK1:Ub using colab notebook [27] and found *h*PINK1:Ub shows similar folding of the ordered Ins3 loop to *Phc*PINK1:Ub post-autophosphorylation [23] (Figure 1b, Figure 1 – figure supplement 1d-g). Although the precise interaction pattern of pSer228 with the Ins3 loop of *h*PINK1 shows variation to the equivalent site at pSer202 in P*hc*PINK1 (Figure 1 – figure supplement 1f-g), several key Ins3 residues of *Phc*PINK1 are conserved in *h*PINK1, including Arg302, Gly307 and the PD associated residue Gly309 (Figure 1b). This suggests a highly conserved pSer228 dependent folding pattern of Ins3 with intolerance of any ins/del in this region across species (Figure 1b).

**Figure 1.**
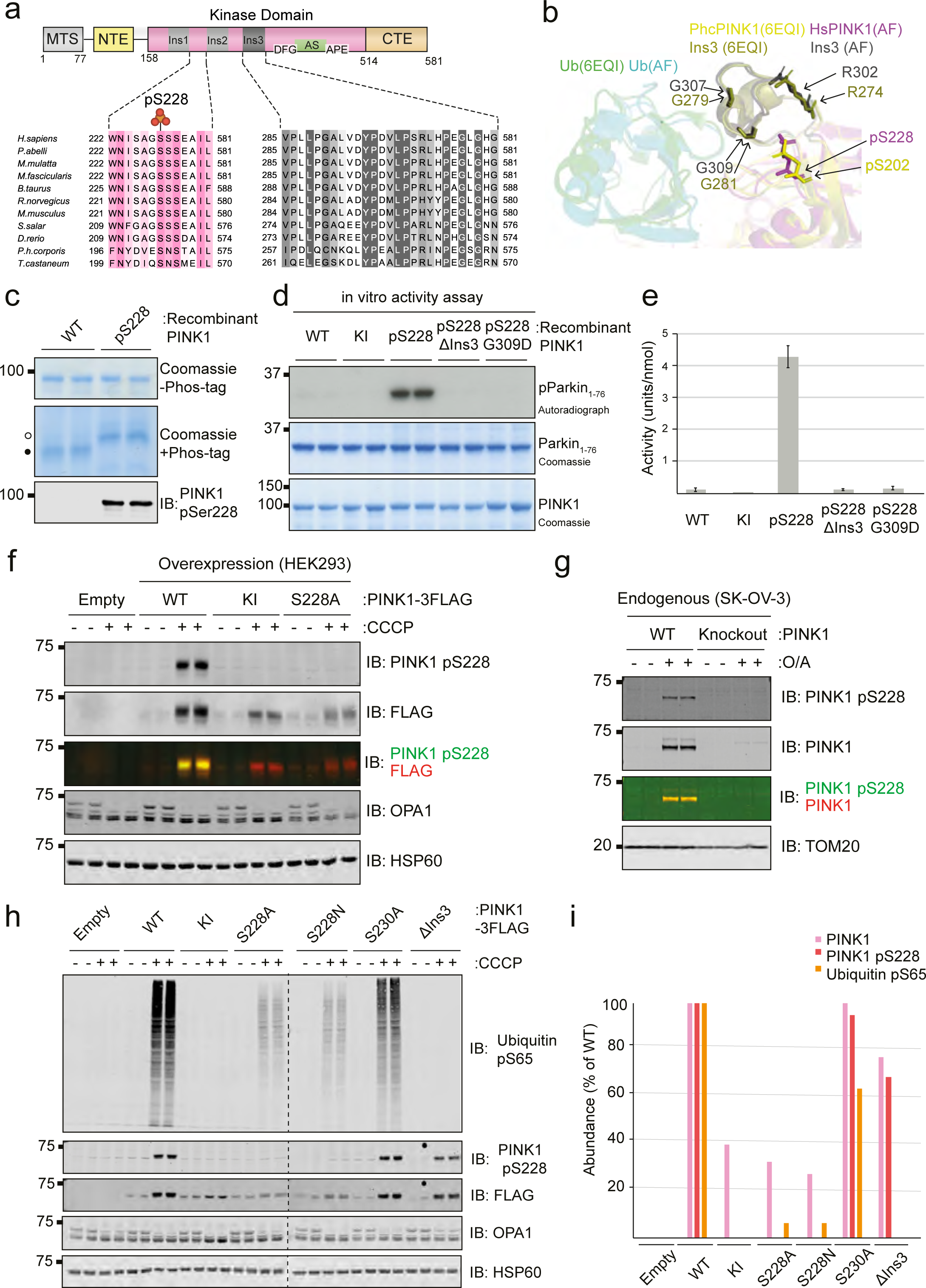
Human PINK1 is activated by phosphorylation at Ser228. **(a)** Conservation of Ser228 and flanking sequences, and Ins3. Sequence alignments were generated with MUSCLE and annotated in JalView. **(b)** Overlapping structures of AlphaFold human PINK1-Ub model and the crystal structure of PhcPINK1 (PDB ID: 6EQI) displaying conserved S228 site and Ins3 residues. (**c)** SDS-PAGE validation of recombinant expression of human PINK1 expressed by normal methods (WT) compared to Ser228-phosphorylated PINK1 generated by genetic code expansion (pS228). 500 ng (for Coomassie staining) or 20 ng (for immunoblotting) were separated by SDS-PAGE, in the presence of 75 μM Phos-tag where indicated. Filled circles indicate non-phosphorylated species; unfilled circles indicate the phosphorylated forms of PINK1. **(d)** *in vitro* kinase assay of recombinant PINK1 comparing activity in presence of absence of pSer228. WT and KI proteins from standard expression conditions were compared to Ser228-phosphorylated PINK1 (pSer228), along with phosphorylated PINK1 further modified by ΔIns3 or incorporation of a pathogenic Ins3 mutation, G309D. Coomassie gels indicate loading of GST-Parkin1-76 substrate and indicated PINK1 proteins. [γ-^32^P] incorporation is visualised by autoradiography following 24 hr exposure. **(e)** Quantitation of PINK1 kinase assay from (d). Gel pieces corresponding to each Parkin1-76 band were excised and analysed by Cerenkov counting. PINK1 activity is expressed as pmol substrate phosphorylated by minute per nmol of kinase (units/nmol). **(f)** Immunoblot validation of pSer228 antibody selectivity. HEK293 cell lines were generated with endogenous PINK1 knocked out and stably replaced by the indicated PINK1-3FLAG mutants. PINK1 expression was induced by 24 hr treatment with 20 nM doxycycline followed by induction of mitochondrial damage by 10 μM CCCP for 3 hr. Mitochondria were harvested and samples analysed by immunoblot (IB), including simultaneous αFLAG / αpSer228 by multiplexing. **(g)** Validation of suitability of pSer228 antibody for detection of endogenous PINK1 autophosphorylation. Mitochondria were harvested from WT and PINK1-knockout SK-OV-3 cells following 9 hr treatment with 10 μM oligomycin and 1 μM antimycin A. Samples were analysed by immunoblotting, including simultaneous αPINK1 / αpSer228 by multiplexing. **(h)** Analysis of PINK1-3FLAG and indicated mutants in HEK293 cell mitochondrial extracts. PINK1-knockout cell lines were generated to stably re-express PINK1-3FLAG WT, kinase inactive (KI; D384A), S228A, S228N, S230A, or with a deletion of residues 295-304 (ΔIns3). PINK1 expression was induced with 20 nM doxycycline for 24 hr, and mitochondrial damage induced by 3 hr treatment with 10 μM CCCP where indicated. Mitochondrial samples were analysed by immunoblotting (IB) with indicated primary antibodies. **(i)** Quantitation of the levels of PINK1, PINK1 pSer228, and ubiquitin pSer65 as in the +CCCP conditions of panel (h). All abundances are expressed as a % of the band intensity relative to WT.

Previous studies have found that recombinant *h*PINK1 expressed in *E. coli* displays near undetectable catalytic activity [20]. We initially explored whether this is due to limited autophosphorylation at Ser228. We observed that recombinant *h*PINK1 resolves as a single band on Phos-tag SDS-PAGE and is not immunoreactive against an anti-phospho-Ser228 PINK1 antibody via immunoblot (Figure 1c). We therefore tested whether artificially phosphorylating recombinant *h*PINK1 at Ser228 would rescue catalytic activity. Employing genetic code expansion technology in an engineered *E. coli* strain [28], we were able to express recombinant hPINK1 (MBP-PINK1_123-581_-6His) stoichiometrically and site-specifically phosphorylated at Ser228 (Figure 1 – figure supplement 2a-e), and confirmed by Phos-tag SDS-PAGE, immunoblotting with an anti-phospho-Ser228 PINK1 antibody (Figure 1c, Figure 1 – figure supplement 2a-d), and mass spectrometry (Figure 1 – figure supplement 2e). Ser228-phosphorylated recombinant *h*PINK1 was also subjected to *in vitro* kinase assay by incubation with recombinant GST-Parkin_1-76_ / Ubl domain, and activity compared to the non-phosphorylated wild-type (WT) and kinase-inactive (KI; D384A) *h*PINK1 proteins expressed in parallel (Figure 1d, e, Figure 1 – figure supplement 2a-e). Excitingly, phosphorylation of Ser228 resulted in a robust increase in *h*PINK1 catalytic activity towards the Parkin substrate (Figure 1d, e, Figure 1 – figure supplement 2f). Consistent with prior structural and *in vitro* analysis of insect PINK1 [21–23], deleting residues 295-304 of Ins3 (ΔIns3) or incorporation of an Ins3 PD pathogenic mutation, G309D, both abolished *h*PINK1 phosphorylation of Parkin even if Ser228 was phosphorylated (Figure 1d, e, Figure 1 – figure supplement 2f). This data confirms that *h*PINK1 utilizes Ins3 as a substrate-binding region, in a manner regulated by autophosphorylation of pSer228. Moreover, these studies confirm that artificially rescuing the autophosphorylation defect of recombinant *h*PINK1 is a tractable method to elaborate *h*PINK1 functional assays *in vitro*.

We deployed the anti-phospho-Ser228 PINK1 antibody to determine whether endogenous *h*PINK1 is phosphorylated at Ser228 in cells. We confirmed the specificity of this antibody using Flp-In T-REx HEK293 PINK1-knockout cells (generated by exon 2-targeted CRISPR-Cas9) re-expressing WT PINK1-3FLAG, KI PINK1 or the non-phosphorylatable S228A mutant by immunoblotting (Figure 1f) and immunofluorescence analysis (Figure 1 – figure supplement 3a). Human wild-type (WT) SK-OV-3 ovarian cell lines or *h*PINK1-knockout (KO) SK-OV-3 cells [26] were treated with or without 10 μM AntimycinA / 1 μM Oligomycin for 9 h to induce mitochondrial depolarisation and immunoblotting of mitochondrial-enriched fractions revealed a specific band for pSer228 that co-migrated with the total *h*PINK1 band in WT cells when the two antibodies were multiplexed, but was abolished in *h*PINK1 KO cells (Figure 1g). These studies provide the first evidence that endogenous *h*PINK1 is capable of autophosphorylation at Ser228 and suggests that monitoring Ser228 autophosphorylation may serve as a robust biomarker of *h*PINK1 catalytic activity. Notably there is 100% sequence identity surrounding PINK1 Ser228 in human, mouse, and rat indicating the potential utility of this antibody in respective PD animal models (Figure 1 – figure supplement 3b).

To determine the role of pSer228 in *h*PINK1 activation in cells, we first generated stable cell lines for expression of PINK1-3FLAG S228A, S228N, or ΔIns3 in Flp-In T-REx HEK293 PINK1-knockout cells and determined pathway activity following CCCP treatment by immunoblotting of mitochondria-enriched lysates (Figure 1h, i). Consistent with our previous analysis in insect PINK1 [21–23], the ΔIns3 mutant prevented ubiquitin recognition and phosphorylation, but preserved Ser228 (trans)autophosphorylation (Figure 1h, i). Ser230 site is significantly phosphorylated in insect PINK1 (*Tc*PINK1 Ser207; *Phc*PINK1 Ser204) but does not contribute to function [21–23] and consistent with this we saw no effect of the S230A mutant on *h*PINK1. Quantification of total *h*PINK1 levels from the mitochondria following CCCP treatment revealed reduced expression levels of the inactive PINK1 mutants (KI, S228A, S228N) (Figure 1h, i). Both S228A and S228N *h*PINK1 mutations drastically reduced ubiquitin phosphorylation at Ser65 following CCCP treatment but in contrast to insect PINK1 analysis [21–23] or the *h*PINK1 KI mutant, there remained minor ubiquitin activity, potentially indicating additional determinants of *h*PINK1 activity (Figure 1h, i).

### Mass Spectrometry identification of *h*PINK1 Ser/Thr phosphorylation sites in cells following mitochondrial depolarisation

We next assessed *h*PINK1 phosphorylation by treating Flp-In T-REx HEK293 *h*PINK1-knockout cells re-expressing WT or KI *h*PINK1-3FLAG with DMSO or CCCP, and then subjecting mitochondria-enriched lysates to Phos-tag SDS-PAGE (Figure 2a, Figure 2 – figure supplement 1a). WT *h*PINK1 exhibits several electrophoretic band shifts on an anti-FLAG immunoblot, indicating multiple phosphorylation sites (Figure 2a, Figure 2 – figure supplement 1a); however, KI *h*PINK1 resolves as one, non-phosphorylated band, strongly suggesting that these modifications of WT *h*PINK1 are due to autophosphorylation (Figure 2a).

**Figure 2.**
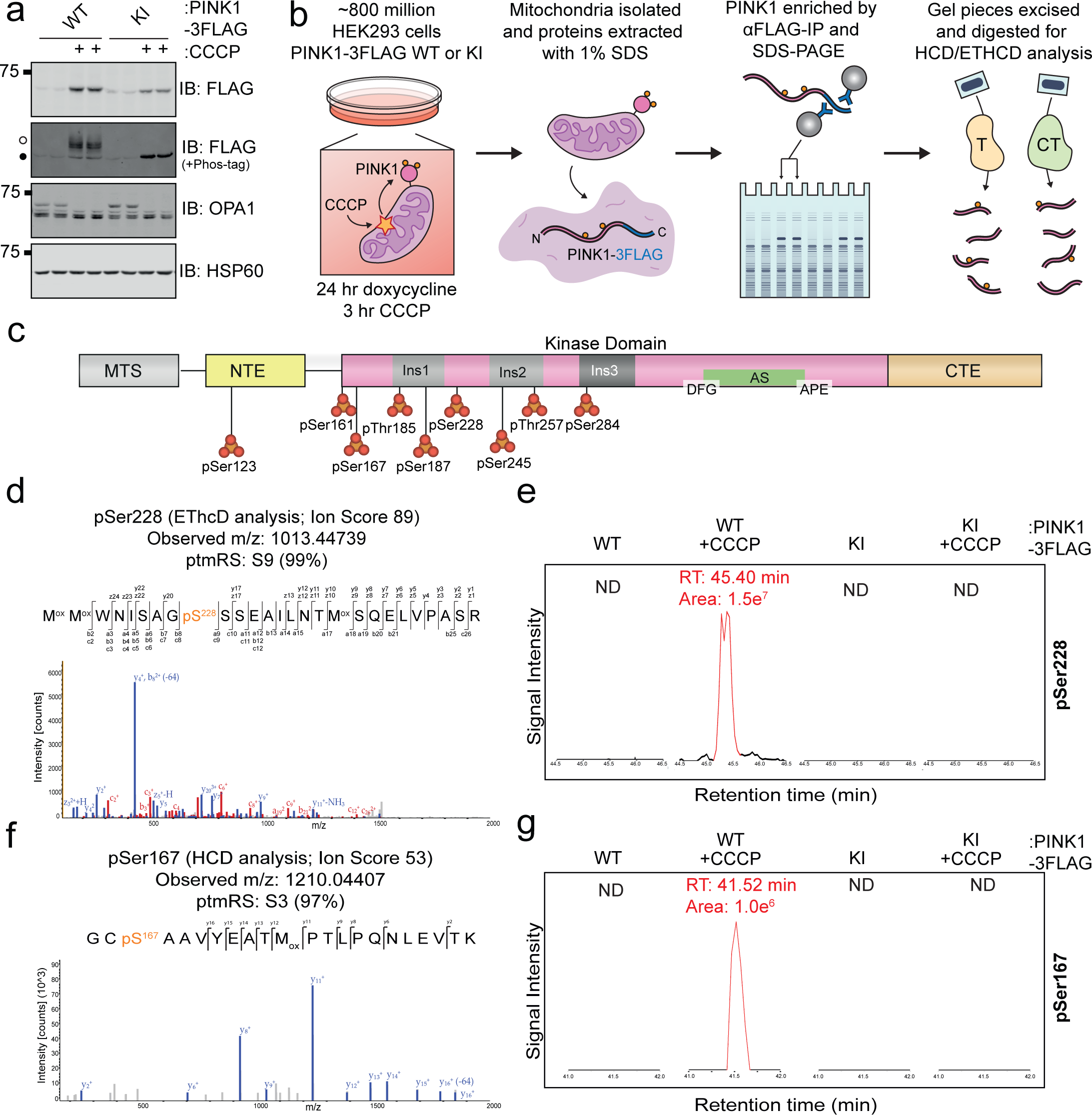
Mass spectrometry-based mapping of human PINK1 Ser/Thr phosphosites. **(a)** Detection of human PINK1 autophosphorylation by Phos-tag SDS-PAGE. PINK1-knockout HEK293 cells with stably re-expressed PINK1-3FLAG as indicated were treated for 3 hrs with CCCP, mitochondria-rich fractions isolated, and samples subjected to SDS-PAGE and immunoblotting (IB). Where specified, SDS-PAGE gels were supplemented with 75 μM Phos-tag reagent for visualisation of phosphorylated species. Filled circles indicate the non-phosphorylated species; unfilled circles indicate the band-shifted and therefore phosphorylated forms. **(b)** Workflow of sample preparation for human PINK1 phosphoproteomics. HEK293 cells from (a) were expanded to ∼800 million cells, treated with doxycycline to induce stable PINK1 expression, and then mitochondria damaged by 3 hr treatment with CCCP. Mitochondria were isolated by differential centrifugation and lysed in 1% buffer to solubilize membrane proteins. PINK1 was immunoprecipitated with αFLAG-amylose resin, then resolved on 4-12% gradient acrylamide gels by SDS-PAGE in duplicate. Gel pieces were excised and subjected to in-gel digestion with either trypsin or chymotrypsin for LC-MS/MS analysis. **(c)** Domain architecture of hPINK1 with mitochondrial targeting sequence (MTS), N-terminal extension (NTE), activation segment (AS), C-terminal extension (CTE), and all identified Ser/Thr autophosphorylation sites indicated. **(d)** Tandem mass spectrum of the triply charged ion at m/z 1013.44739 following EThcD fragmentation with a Mascot ion score of 89 and ptmRS phosphosite localisation confidence of 99% indicating phosphorylation at Ser228. **(e)** Label free quantification of pSer228 peptide across treatment conditions. Extracted ion chromatograms (XICs) were performed on the first isotopic peak (*m/z* 1013.7818), which had the highest signal intensity. No precursor with *m/z* 1013.7818 was detected in any condition other than WT + CCCP. Peaks with the correct *m/z* and retention time are highlighted in red. RT: retention time. (**f)** Tandem mass spectrum of the doubly charged ion at m/z 1210.04407 following HCD fragmentation with a Mascot ion score of 53 and a ptmRS phosphosite localisation confidence of 97% indicating phosphorylation at Ser167. **(g)** Label free quantification of pSer167 peptide across treatment conditions. XICs were performed on the monoisotopic peak (*m/z* 1210.0441). No precursor with *m/z* 1210.0441 was detected in any condition other than WT + CCCP. Peaks with the correct m/z and retention time are highlighted in red. RT: retention time.

To map the *h*PINK1 phosphorylation sites using an unbiased approach, we prepared samples for mass spectrometry (MS) analysis. Using MS, we previously reported a non-regulatory Threonine257 (Thr257) autophosphorylation site [13], however, other sites were not robustly detected including Ser228 [13]. This contrasted with multiple-phosphorylation events observed on Phos-tag SDS-PAGE (Figure 2a, Figure 2 – figure supplement 1a) leading us to optimise mitochondrial lysis conditions to maximise isolation of cellular *h*PINK1. We have previously employed lysis buffers containing 1% Triton X-100, and we observed that this failed to fully solubilize mitochondrial *h*PINK1 (Figure 2 – figure supplement 1b, c). Furthermore, immunoblotting of Triton X-100 soluble and insoluble fractions revealed that Ser228-phosphorylated *h*PINK1 was predominantly in the Triton-insoluble fraction (Figure 2 – figure supplement 1d) which explains the lack of robust detection of Ser228 phosphorylation in previous MS analysis [13], In contrast, lysis in 1% SDS maximally solubilizes mitochondrial *h*PINK1, and was therefore used in subsequent analysis (Figure 2 – figure supplement 1b, c).

Immunoprecipitates of WT or KI *h*PINK1-3FLAG from SDS-solubilised mitochondrial extracts (from CCCP-treated and untreated HEK293 cells) were separated by SDS-PAGE, and gel pieces corresponding to ∼63 kDa excised for in-gel proteolytic digestion with either trypsin or chymotrypsin (Figure 2b, Figure 2 – figure supplement 1e). The resulting peptides were analysed by liquid chromatography (LC)-tandem MS (MS/MS) using either Higher energy Collisional Dissociation (HCD) or Electron-transfer/HCD (EThcD) fragmentation techniques (Figure 2b). A total of 8 *h*PINK1 Ser/Thr phosphorylation sites were unambiguously identified, all exclusively present on WT *h*PINK1 following CCCP treatment (Figure 2c-g, Figure 2 – figure supplements 2 and 3). Trypsin (Figure 2 – figure supplement 2a) and Chymotrypsin (Figure 2 – figure supplement 3a) digests gave a combined sequence of ∼75% across *h*PINK1 (Figure 2 – figure supplement 3c).

LC-MS/MS analysis with EThcD fragmentation revealed a tryptic peptide containing phosphorylated Ser228 in WT *h*PINK1-3FLAG cell lines treated with CCCP (Figure 2d); however, this modification was absent in untreated WT cells, or in cells expressing *h*PINK1-3FLAG KI consistent with this being an autophosphorylation site (Figure 2d, e). Previous studies have been unable to detect Ser228 phosphorylation by MS since it is located in a triple-serine motif, Ser^228^Ser^229^Ser^230^ [29]. To our knowledge, the peptide detected here represents the first unambiguous MS detection of Ser228 phosphorylation in *h*PINK1, with fragmentation patterns site-specifically pinpointing phosphorylation on Ser228 over the other serine residues, based in part on the observation of b_8_/c_8_ and a_9_/c_9_ product ions (Figure 2d). Notably, phosphorylation of Ser230 could not be detected, despite this residue being well conserved and robustly phosphorylated on insect PINK1 [20–23].

### Site conservation and structural evaluation of *h*PINK1 Ser/Thr phosphorylation sites

Our analysis unambiguously identified six novel Ser/Thr autophosphorylation sites (pSer123, pSer161, pSer167, pThr185, pSer187, and pSer245) specific to wild-type *h*PINK1 (Figure 2 c, f, g; Figure 2 – figure supplements 2 and 3); pThr257 (Figure 2 – figure supplement 2f) [13] and a phosphorylated peptide containing the previously reported pSer284 autophosphorylation site [30] (data not shown). We next assessed conservation at these sites (Figure 2 – figure supplement 4) and modelled their location utilising the *h*PINK1-ubiquitin complex structure model from the EMBL-EBI AlphaFold2 (AF2) database (AFDB) (Figure 2 – figure supplement 5) [27]. pSer167 (Figure 2f, g and Figure 3a) lies in a commonly phosphorylated location in the kinase glycine-rich loop [31], however, inspection of the *h*PINK1-ubiquitin complex model indicated that pSer167 also lies at the interface with ubiquitin (Figure 2 – figure supplement 5b). Moreover, according to the AF2 *h*PINK1: Ub model, there is adequate space at this interface to facilitate Ser167 phosphorylation (Figure 3b).

**Figure 3.**
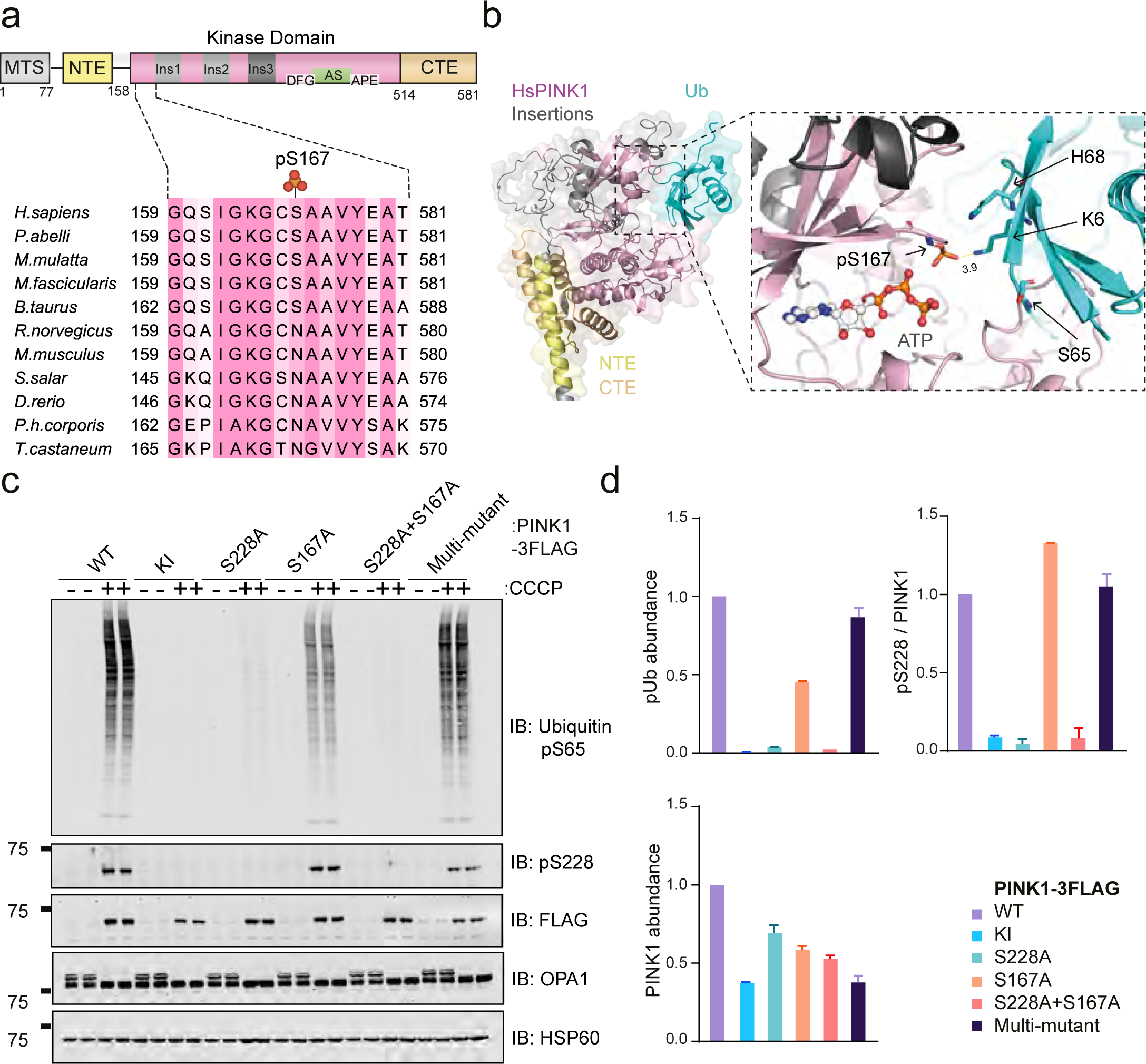
Ser167 phosphorylation is required for efficient Human PINK1 ubiquitin phosphorylation. **(a)** Conservation of Ser167 site. Sequence alignments were generated with MUSCLE and annotated in JalView. **(b)** AlphaFold human PINK1-Ub model showing interaction of pS167 with positively charged residues of Ubiquitin **(c)** Analysis of PINK1-3FLAG and indicated mutants in Hela cell mitochondrial extracts. PINK1-knockout cell lines were generated to stably re-express PINK1-3FLAG WT, kinase inactive (KI; D384A), S228A, S167A, S228A + S167A or with a multi-mutant (T185A, S187A, T257A, S284A, S73A, S123A, S245A). PINK1 expression was induced with 20 nM doxycycline for 24 hr, and mitochondrial damage induced by 3 hr treatment with 10 μM CCCP where indicated. Mitochondrial samples were analysed by immunoblotting (IB) with indicated primary antibodies. **(d)** Quantitation of the levels of PINK1, PINK1 pSer228, and ubiquitin pSer65 as in the +CCCP conditions of panel (c). All abundances are expressed as the band intensity relative to WT.

Furthermore, Ub presents two positively charged residues, Lys6 and His68, located in close proximity to the negatively charged pSer167 (Figure 3b). The formation of a salt-bridge interaction between Lys6 and pSer167 implies a potential role of pSer167 in facilitating interaction and binding of Ub (Figure 3b).

pThr185, pSer187, pSer245, and the known pThr257 site [13] are located within *h*PINK1’s first two kinase domain insertions (Ins1, Ins2) although structural prediction of Ins1 and Ins2 is limited by low confidence of the AF2 model in this region (Figure 2c, Figure 2 – figure supplements 5 a-f). We have previously found that that these insertions can be deleted in *h*PINK1 with negligible effect on *h*PINK1 activity in cells [21]. The previously reported pSer284 [30] is predicted to be located within Ins3 (Figure 2c, Figure 2 – figure supplements 5 a, b, g, h). and pSer123 is located within the N-terminal extension (NTE) (Figure 2c, Figure 2 – figure supplements 5 a, b, g, h) whose structural predictions are high in the model. Ser161 is located within the kinase glycine-rich loop but away from the ubiquitin interface (Figure 2c, Figure 2 – figure supplements 5 a, b, i, j). Interestingly apart from Ser228, these other phosphorylated Ser/Thr residues are not conserved in the previously characterised insect orthologues, *Tc*PINK1 and *Phc*PINK1 (Figure 2 – figure supplement 4).

To address the functional role of all phosphorylation sites that we mapped, we generated Flp-In T-REx HeLa *h*PINK1-knockout cells stably re-expressing WT *h*PINK1-3FLAG or mutant KI, S228A, S167A, S228A/S167A or a combination mutant of S/T residues not at the ubiquitin interface including S123A/S245A/T185A/S187A/T257A/S284A (multi-mutant). Cells were treated with DMSO or CCCP followed by immunoblotting of mitochondrial-enriched lysates (Figure 3c). Consistent with the AF2 structural model of *h*PINK1-ubiquitin, we observed that the S167A mutant significantly inhibited ubiquitin phosphorylation, but preserved Ser228 (trans)autophosphorylation and furthermore the residual ubiquitin phosphorylation of the S228A mutant was further reduced in the S167A/S228A double mutant (Figure 3c-d). In contrast we observed a moderate effect of the multi-mutant on *h*PINK1 stabilisation, but this did not significantly impact ubiquitin phosphorylation or Ser228 (trans)autophosphorylation (Figure 3c). These analyses indicate that *h*PINK1 catalysed ubiquitin phosphorylation is dependent on pSer167 in addition to pSer228. Furthermore, the Ins2 phosphorylation site, Thr257, is dispensable for *h*PINK1 activation and Ser228 (trans)autophosphorylation in marked contrast to its role in the active dimeric *Tc*PINK1 structure [25].

### Identification and functional analysis of novel *h*PINK1 Cys412 phosphorylation

In agreement with previous work investigating non-canonical phosphorylation *e.g.* of cysteine in human cells [32], we detected phosphorylation of Cys412 following EThcD analysis in WT *h*PINK1-3FLAG cells (Figure 2 – figure supplement 2 g, h). However, this peptide was not consistently detected in independent replicate EThcD experiments (data not shown) suggesting that this modification is labile and not stable under this MS fragmentation method.

Collision-mediated fragmentation strategies such as HCD and collision-induced dissociation (CID) resulting in proton-directed bond cleavage is known to be problematic for the interrogation of labile post-translational modifications (PTMs). In contrast, electron-driven fragmentation strategies are better at retaining labile PTMs and thus permitting site localisation [33]. We therefore explored whether the pCys412 *h*PINK1 was detectable by Electron Activated Dissociation (EAD) using a ZenoTOF7600 mass spectrometer that enables tuneable and rapid EAD [34]. Samples were prepared from immunoprecipitates of WT (-/+ CCCP), or KI (+CCCP) *h*PINK1-3FLAG from SDS-solubilised mitochondria extracts (from CCCP-treated and untreated stable HEK293 cells); separated by SDS-PAGE, and gel pieces corresponding to ∼63 kDa excised for in-gel proteolytic digestion with trypsin/LysC mix (Figure 4 – figure supplement 1a-c). The resultant peptides were analysed by EAD or CID without phosphopeptide enrichment in a blinded fashion and revealed the phosphorylated peptide encompassing Cys412 was detected using EAD but not with CID (Figure 4a, Figure 4 – figure supplement 1d). WT *h*PINK1 + CCCP is stabilised ∼10fold more than WT *h*PINK1 – CCCP or the KI PINK1 + CCCP (Figure 4 – figure supplement 1b, c) and we next employed Multiple Reaction Monitoring (MRM) to quantify the peptide containing pCys412 in each cell sample relative to FLAG peptide (Figure 4b). This revealed that pCys412 is present constitutively in WT - *h*PINK1 and KI+ *h*PINK1 and is increased in WT + *h*PINK1 in a manner that correlates with peptide abundance (Figure 4b).

**Figure 4.**
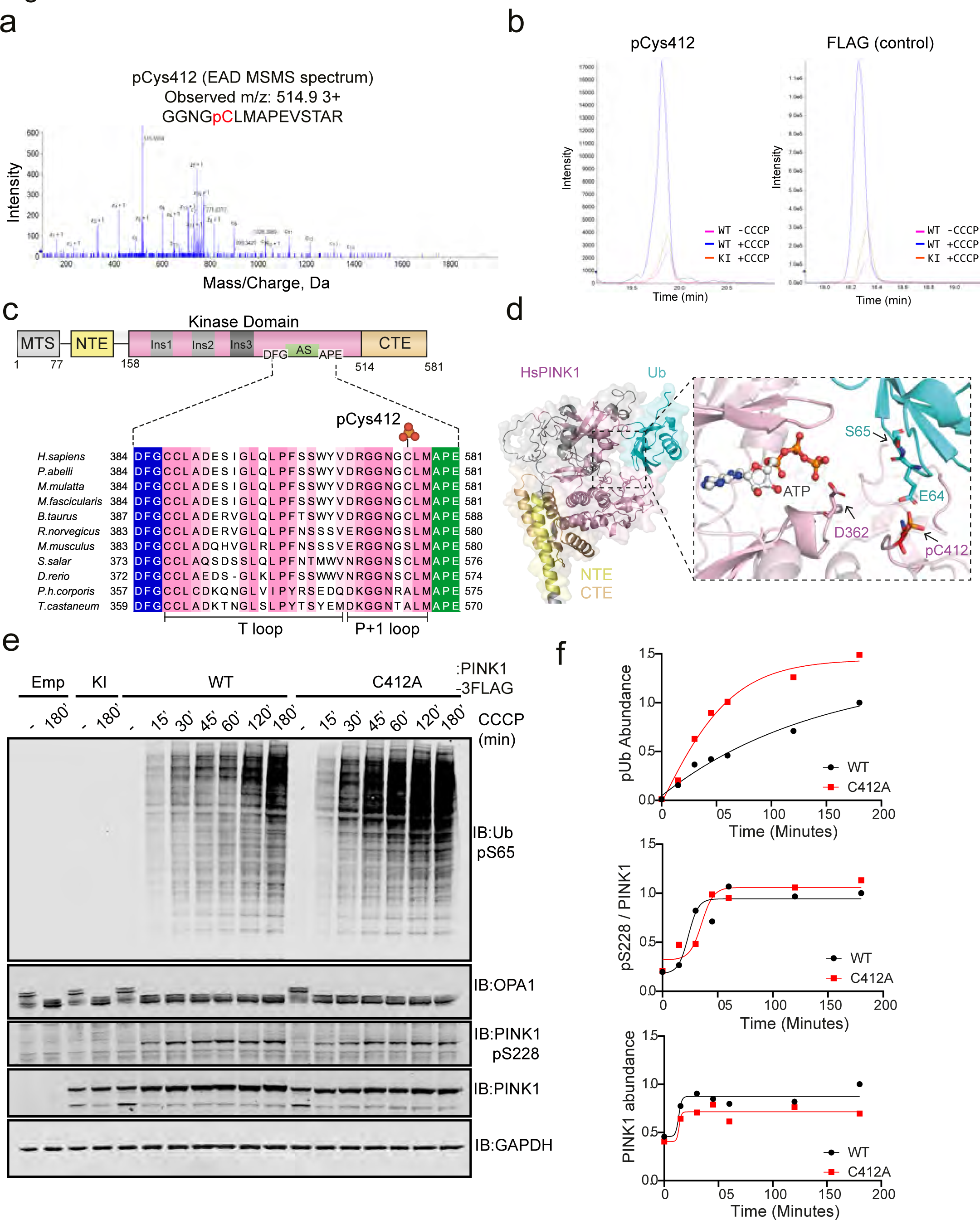
Human PINK1 exhibits phosphorylation at Cys412. **(a)** EAD MS/MS spectrum of GGNGpCLMAPEVSTAR (*m/z* 514.9 3+). All matched c and z+1 ions are annotated in the spectrum. **(b)** (Left) Targeted MRMhr experiments was performed on peptide GGNGpCLMAPEVSTAR (*m/z* 514.9 3+) using optimised collision energies. The extracted ion chromatograms are for the WT PINK + CCCP (blue), WT PINK-CCCP (pink), KI PINK (orange). The extracted ion chromatogram for GGNGpCLMAPEVSTAR (*m/z* 514.9 3+) is the sum of the fragment ions *m/z* 830,437, 759.399, 662.347, 533.304 and 434.236 representing the y8-y4 ions for this peptide extracted with a tolerance of +/- 0.01Da. (Right) Targeted MRMhr experiments was performed on peptide AALDYMDDDDK (*m/z*636.3 2+) using optimised collision energies. The extracted ion chromatograms are for the WT PINK + CCCP (blue), WT PINK-CCCP (pink) and KI PINK (orange). The extracted ion chromatogram for AALDYMDDDDK (*m/z* 636.3 2+) is the sum of the fragment ions *m/z* 738.261, 607.220, 492.193 and 377.167 representing the y6-y3 ions for this peptide extracted with a tolerance of +/- 0.01Da. **(c)** Conservation of PINK1’s activation segment across species. Sequence alignments were generated with MUSCLE and annotated in JalView. Regions corresponding to the T loop and P+1 loop are indicated below. **(d)** AlphaFold model of hPINK1 and Ubiquitin. ATP is superimposed into the hPINK1 active site based on its position in the complex structure. **(e)** Time-course analysis of PINK1-3FLAG and indicated mutants in HEK293 whole cell lysates. PINK1-knockout cell lines were generated to stably re-express PINK1-3FLAG WT, kinase inactive (KI; D384A), empty-FLAG and C412A. PINK1 expression was induced with 20 nM doxycycline for 24 hr, and mitochondrial damage induced by 3 hr treatment with 10 μM CCCP where indicated. Samples were analysed by immunoblotting (IB) with indicated primary antibodies. **(f)** Quantitation of the levels of PINK1, pSer228, and ubiquitin Ser65 as in the +CCCP conditions of panel (c). All abundances are expressed as a band intensity relative to wildtype.

Cys residues play important emerging regulatory roles in the activation segments of eukaryotic Ser/Thr kinases [35]. Cys412 is highly conserved in higher eukaryotes, although it is replaced by a serine in rodents (Figure 4c). Notably, however, Cys412 is absent in insect species, where it is replaced by an alanine in both *Tc*PINK1 and *Phc*PINK1 (Figure 4c). Cys412 maps to the P+1 loop – a subregion at the end of the activation segment that determines substrate specificity by recognising the residues flanking the substrate phosphorylation site. To evaluate possible functional effects of Cys412 phosphorylation, we inspected the active site in the AF2 *h*PINK1-ubiquitin complex structure model (Figure 4d). This predicts that pCys412 would be incompatible with ubiquitin binding, either occupying space or electrostatically repelling the side chain of Glu64 of the ubiquitin substrate. This suggests that P+1 loop phosphorylation at Cys412 would inhibit *h*PINK1 by preventing the binding of substrate. To test this hypothesis, we generated stable cell lines for the expression of *h*PINK1-3FLAG C412A in the PINK1-knockout HEK293 background. Strikingly we observed that prevention of phosphorylation with a C412A mutation increased ubiquitin phosphorylation, in agreement with a possible role for pCys412 in preventing substrate recognition (Figure 4e-f). Consistent with this, we did not observe any significant change in levels of PINK1 C412A or (trans)autophosphorylation of Ser228 (Figure 4e-f).

## Conclusion

The activation mechanism of *h*PINK1 is of great biomedical interest, since small molecule activators of *h*PINK1 have significant potential as a therapeutic strategy against Parkinson’s disease (PD). We demonstrate that the weak phosphotransferase activity of recombinant *h*PINK1 *in vitro* is due to an inability to autophosphorylate at Ser228, and that by directly incorporating phosphoserine during recombinant expression, *h*PINK1 activity can be rescued (Figure 1). Using this reconstituted system, we provide direct evidence that active *h*PINK1 utilises Ins3 as a substrate-binding loop controlled by pSer228 (Figure 1) as predicted by previous structural studies of insect PINK1 orthologues [21–23]. Next, we undertook the most comprehensive MS-based analysis of *h*PINK1 Ser/Thr phosphorylation leading to the identification of pSer167 as a regulatory autophosphorylation site of *h*PINK1 in addition to pSer228 (Figures 2 and 3) and the discovery of six novel biomarker sites of *h*PINK1 activity (Figures 2 and 3). Strikingly we also identified phosphorylation of a conserved Cys412 within the activation segment of *h*PINK1 and functional studies indicate a potential inhibitory role (Figure 4). We observe that Cys412 phosphorylation is constitutive, being present on WT *h*PINK1 under basal conditions and KI *h*PINK1. Structural modelling indicates that pCys412 limits accessibility of Ubiquitin Ser65 into the active site for phosphorylation by *h*PINK1 catalytic machinery (Figure 4d). Under basal conditions *h*PINK1 undergoes proteasomal degradation and we hypothesise that pCys412 would provide another mechanism to ensure *h*PINK1 is kept inactive to prevent inappropriate induction of mitophagy (Figure 4 figure supplement 2). Upon mitochondrial damage we postulate that the stoichiometry of autophosphorylated *h*PINK1 is greater than pCys412 *h*PINK1 resulting in overall activation (Figure 4 figure supplement 2). In eukaryotes cysteine phosphorylation is known to be important as a catalytic intermediate in the function of protein tyrosine phosphatases (PTPs) [36]. Our data suggests that pCys may also have a role in directly regulating eukaryotic cell signalling similar to a previous observation In prokaryotes, where proteins of the SarA and MgrA family of transcription factors of *Staph aureus* were demonstrated by mass spectrometry to be phosphorylated at Cysteine residues mediated by the phosphatase Stp1 and kinase Stk1 [37]. In future work it would be interesting to uncover the mechanism of *h*PINK1 Cys412 phosphorylation and determine whether this is an enzyme-mediated process driven *e.g.* by a mitochondrial homologue of a prokaryotic cysteine kinase or arises from phosphate transfer linked to high reactivity of the Cys thiol group. To our knowledge, this is the first example of a protein kinase regulated by cysteine site phosphorylation, as opposed to cysteine oxidation [35] in the activation segment. Whole-kinome alignments reveal that the CxxAPE motif found in the *h*PINK1 P+1 loop is also present in OSR1, NEK8, STK39, KSR1 and KSR2 [35]. Since the APE motif lies in a very similar structural position in all protein kinases, it would be interesting to determine whether these −3 position cysteines are also phosphorylated similar to *h*PINK1 Cys412 and if so, assess how modification impacts on substrate binding and catalytic activity. Overall, our findings provide new insights into *h*PINK1 activation and may aid in the development of new biomarkers and strategies to enhance *h*PINK1 activation.

## Methods

### Reagents and cloning

[γ-^32^P] ATP was obtained from PerkinElmer. *Hs*PINK1-3FLAG constructs were cloned into pcDNA5 vectors for recombination into HEK293 Flp-In TRex/Hela cell lines. MBP-HsPINK1 constructs were cloned into pMal4C vectors for recombinant protein expression, whereas MBP-HsPINK1-6His constructs for genetic code expansion were cloned into pNHD1.3 vectors (17). Site-directed mutagenesis was carried out using the QuikChange method with KOD polymerase (Novagen). All constructs were verified by The Sequencing Services (School of Life Sciences, University of Dundee) and are now available to request via the MRC PPU Reagents and Services website (https://mrcppureagents.dundee.ac.uk/).

### Antibodies

The following antibodies were used in this study: α-FLAG (Sigma), α-HSP60 (CST), α-OPA1 (BD biosciences, 612606), α-PINK1 (Novus, BC100-494), α-PINK1 (DCP), α-ACSL1 (Cell signalling technologies, 4047S), α-ubiquitin pSer65 (Cell signalling technologies, 37642). The polyclonal α-PINK1 pSer228 antibody was generated by the Michael J. Fox Foundation’s research tools program in partnership with Abcam (Development of a monoclonal antibody is underway. Please contact for questions.) All fluorophore-conjugated mouse, rabbit, and sheep secondary antibodies for immunoblotting and immunofluorescence were obtained from Sigma.

### Mammalian cell culture

SK-OV-3 and HeLa wild-type (WT) and PINK1 knockout cells were routinely cultured in standard DMEM (Dulbecco’s modified Eagle’s medium) supplemented with 10% FBS (fetal bovine serum), 2 mM L-Glutamine, 100 U ml−1 Penicillin, 100 mg ml−1 Streptomycin (1X Pen/Strep) and 1 X non-essential amino acids (Life Technologies). HEK293 Flp-In TREx cells/Hela Flp-In cells were cultured using DMEM (Dulbecco’s modified Eagle’s medium) supplemented with 10% FBS (foetal bovine serum), 2 mM L-glutamine, 1x Pen/Strep, and 15 μg/ml blasticidin. Culture media was further supplemented with 100 μg/ml zeocin pre-recombination with PINK1-3FLAG constructs. Transfections were performed using the polyethylenimine method. To ensure uniform expression of recombinant proteins, stable cell lines were generated in a doxycycline-inducible manner. Flp-In T-REx-293/Flp-In T-rex-Hela CRISPR knockout to PINK1 null cells were generated in a laboratory. The PINK1 null host cells containing integrated FRT recombination site sequences and Tet repressor were co-transfected with 4.5/9 µg of pOG44 plasmid (which constitutively expresses the Flp recombinase) and 0.5/1 µg of pcDNA5-FRT/TO vector containing a hygromycin resistance gene for selection of the gene of interest with FLAG tag under the control of a doxycycline-regulated promoter. Cells were selected for hygromycin and blasticidin resistance 3 days after transfection by adding fresh medium supplemented with 15 µg ml−1 of blasticidin and 100 µg ml−1 of hygromycin. Protein expression was induced by the addition of 0.1 μg/ml doxycycline for 24 hours. Mitochondrial depolarisation was induced by treatment with 10 μM CCCP (Sigma; prepared in DMSO) for 3 hrs.

Cells were harvested and resuspended in mitochondrial fractionation buffer (20mM HEPES pH 7.5, 250 mM sucrose, 3 mM EDTA, 5 mM sodium β-glycerophosphate, 50 mM sodium fluoride, 5 mM sodium pyrophosphate, 1 mM sodium orthovanadate, 200 mM chloracetamide). Cell suspensions were physically disrupted by 25 passes through a 25-gauge needle, and debris removed by centrifugation at 800 x g. The resulting supernatant was subject to centrifugation at 16,600 x g to harvest a mitochondria-enriched pellet. Unless indicated, samples for SDS-PAGE were generated from mitochondrial lysates resulting from resuspension in mitochondrial fractionation buffer with 1% SDS.

### Immunoprecipitation

PINK1-3FLAG was enriched from HEK293 stable cell lines by α-FLAG-immunoprecipitation. Mitochondria were isolated by differential centrifugation and solubilised in mitochondrial fractionation buffer with 1% (w/v) SDS. Mitochondrial lysates were incubated for 16hrs with αFLAG-agarose (200:1 ratio; MRC PPU Reagents and Services), in a volume sufficient to reduce the final SDS concentration to 0.1% (w/v). Supernatants were retained to determine immunodepletion. Non-specifically bound proteins were removed by three washes in mitochondrial fractionation buffer with 150 mM NaCl, followed by one wash in mitochondrial fractionation buffer to re-equilibrate. Proteins were eluted by the addition of 1x LDS and incubation at 70°C with shaking for 15 mins. Denatured eluates were separated from the beads by centrifugation through a 0.22 μM Spin-X column. For phosphoproteomic analysis, samples were reduced by the addition of β-mercaptoethanol to a final concentration of 1% (v/v). During optimisation steps, a further quality control step was performed by direct addition of 1x LDS with 1% (v/v) β-mercaptoethanol to the isolated beads and re-incubation at 70^ο^ C, therefore enabling visualisation and quantification of proteins not removed by the initial elution step (“uneluted”). IP efficiency was determined by SDS-PAGE and α-FLAG immunoblot with normalised loading of input (mitochondrial lysate), supernatant, and eluate samples.

### Sample preparation for phosphoproteomic analysis

The phosphorylation status of human PINK1-3FLAG (immunoprecipitated from 20 mg of SDS-lysed HEK293 cells mitochondria) was probed by phosphoproteomics. Samples were reduced in 10 mM DTT for 10 mins at 90 °C, then allowed to cool prior to 30 mins alkylation with 20 mM iodoacetamide. Alkylated samples were analysed by SDS-PAGE on 4-12% acrylamide gradient gels. Gels were stained in colloidal Coomassie, and bands corresponding to the molecular weight of PINK1-3FLAG (∼63 kDa) were excised and cut into 1 mM cubic pieces. Gel pieces were dehydrated with 100% acetonitrile, then rehydrated with 50 mM ammonium bicarbonate three times. Gel pieces were then dried down via vacuum centrifugation, then rehydrated in 25 mM triethylammonium bicarbonate (TEAB) containing either 1 μg trypsin, chymotrypsin, or Asp-N proteases where indicated for overnight digestion. The following day, gel pieces were re-shrunk by the addition of an equal volume of acetonitrile, and the resulting peptide-containing 50% acetonitrile, 12.5 mM TEAB solution removed and dried by Speedvac. Dried samples were resuspended in 0.5% (v/v) trifluoracetic acid and desalted with C18 spin tips, then dried again by Speedvac for storage at −20 °C. Final dried peptides were resuspended in 0.1% (v/v) formic acid immediately before LC-MS/MS analysis.

### Phosphoproteomic analysis by liquid chromatography mass spectrometry (LC-MS/MS)

LC separations were performed with a Thermo Dionex Ultimate 3000 RSLC Nano liquid chromatography instrument using 0.1% formic acid as buffer A and 80% acetonitrile with 0.08% formic acid as buffer B. The peptide samples were loaded on C18 trap columns with 3% acetonitrile / 0.1% trifluoracetic acid at a flow rate of 5 μL/min. Peptide separations were performed over EASY-Spray column (C18, 2 μM, 75 μm x 50 cm) with an integrated nano electrospray emitter at a flow rate of 300 nL/min. Peptides were separated with a 60 min segmented gradient starting from 5%∼30% buffer B for 30 min, 30%∼45% buffer B for 15 mins, 45%∼95% buffer B for 5 min, followed by 95% buffer B for 5 min. Eluted peptides were analysed on an Orbitrap Fusion Lumos (ThermoFisher Scientific, San Jose, CA) mass spectrometer. Spray voltage was set to 2 kV, RF lens level was set at 30%, and ion transfer tube temperature was set to 275 °C. The Orbitrap Fusion Lumos was operated in positive ion data dependent mode. The mass spectrometer was operated in data-dependent top speed mode with 3 seconds per cycle. The full scan was performed in the range of 375—1500 *m/z* at nominal resolution of 120,000 at 200 *m/z* and AGC set to 4 x10^5^ with maximal injection time of 50 ms, followed by selection of the most intense ions above an intensity threshold of 5000 for electron-transfer/higher-energy collision dissociation (EThcD) fragmentation (18,19). EThcD spectra were collected using calibrated charge-dependent ETD parameters and HCD supplementary activation was enabled. HCD normalised collision energy was set to 30% for EThcD. The anion AGC target was set to 4 x 10^5^ for EThcD. Data-dependent MS2 scans were acquire for charge states 2 to 7 using an isolation width of 1.6 Th and a 20 s dynamic exclusion. All MS2 scans were recorded in the ion trap with centroid mode using an AGC target of 1 x 10^4^ and a maximal fill time of 35 ms. Whereas for HCD, MS2 spectra were recorded in the orbitrap in centroid mode, with nominal resolution of 15,000 at 200 m/z, AGC set to 5×10^4^ and a maximum injection time of 50ms. The .RAW files obtained from the mass spectrometer were analysed by Proteome Discoverer v2.2 (ThermoFisher Scientific) using Mascot v 2.6.2 (Matrix Science) as the search engine. A precursor mass tolerance of 10ppm and fragment tolerance of 0.6 was used. Swissport database (downloaded in April 2020 from Uniprot) was used with taxonomy restriction to *Homo sapiens*. It was allowed a maximum of two missed cleavage sites and the protease was selected according to the samples being analysed (trypsin or chymotrypsin). Oxidation of methionine, phosphorylation of serine, threonine, and cysteine, and carbamidomethylation of cysteine were set as variable modifications. ptmRS was used as a scoring system for the phospho site identification, with a mass tolerance of 0.5 and neutral loss peaks were considered for HCD and EThcD phosphorylation site localisation was considered correct only if peptides had a Mascot score above 28 and ptmRS probability score above 85%.

Cysteine modifications being labile, the samples for EAD analysis of phosphorylated cysteine residues were prepared with slight modifications to the methods for EThcD. Samples were reduced in 10 mM DTT for 20 mins at 30 °C, then allowed to cool prior to 30 mins alkylation with 20 mM iodoacetamide. Alkylated samples were analysed by SDS-PAGE on 4-12% acrylamide gradient gels. Gels were stained in colloidal Coomassie, and bands corresponding to the molecular weight of PINK1-3FLAG (∼63 kDa) were excised and cut into 1 mM cubic pieces. Gel pieces were dehydrated with 100% acetonitrile, then rehydrated with 50 mM ammonium bicarbonate three times. Gel pieces were then dried down using vacuum centrifugation, then rehydrated in 25 mM triethylammonium bicarbonate (TEAB) containing 1 µg Trypsin/Lys-C mix for overnight digestion. The following day, digested peptides were eluted by adding 80% (v/v) acetonitrile in 0.5% formic acid (2 times), 100% acetonitrile (1 time) pooled and were dried by Speedvac. Dried samples were resuspended in 0.5% (v/v) trifluoracetic acid and desalted with C18 spin tips, then dried again by Speedvac for storage at −20 °C. Final dried peptides were resuspended in 0.1% (v/v) formic acid immediately before EAD analysis on ZenoTOF 7600.

Bacterial recombinant hSer228PINK1 digests were separated with a Thermo Dionex Ultimate 3000 RSLC Nano liquid chromatography instrument configured as above with the peptides being separated with a 42 min segmented gradient starting from 3%∼35% buffer B for 40 minutes and 35%∼99% buffer B over 2 minutes. The eluted peptides were analysed on an Orbitrap Velos (ThermoFisher Scientific, San Jose, CA) mass spectrometer and fragmented by collision induced dissociation using a normalised collision energy of 35. The acquired raw files were processed by Proteome Discoverer v2.2 (ThermoFisher Scientific) and the resulting data searched against an inhouse database (MRC_Database_1) using Mascot v 2.6.2 (Matrix Science) search engine. It was allowed a maximum of two missed cleavage sites with a peptide tolerance of 10ppm and a fragment tolerance of 0.6 Daltons. The protease was specified as trypsin. Oxidation and Dioxidation of methionine, phosphorylation of serine, threonine and tyrosine were set as variable modifications. Carbamidomethylation of cysteine was set as a fixed modification.

### PhosTag™ SDS-PAGE

The phosphorylation status of PINK1 was determined by Phos-tag™ SDS-PAGE using a protocol optimised by Okatsu *et al* (10). Tris-Glycine gels were prepared with 7.5% acrylamide, 50μM Phos-tag™ acrylamide AAL-107 (Wako) and 100μM MnCl_2_. In order to improve efficiency of wet transfer, run gels were washed three times in standard transfer buffer with the addition of 0.02% SDS and 5mM EDTA, followed by one wash in transfer buffer with 0.02% SDS only. Similarly, transfer times were extended to 180 mins at 90V to improve transfer of high molecular weight proteins.

### Immunofluorescence

PINK-3FLAG-expressing HEK293 cell lines were seeded onto coverslips, pre-coated with poly-L-Lysine overnight. PINK1 expression was induced by 24 hr treatment with doxycycline, with 3 hr CCCP treatment for mitochondrial uncoupling. Following treatments, coverslips were washed twice in phosphate-buffered saline (PBS), and cells fixed by the addition of 3.7% (v/v) formaldehyde for 20 mins. Remaining formaldehyde was removed by three washes in DMEM + 10 mM HEPES pH 7.0, followed by a further wash in PBS. Coverslips were permeabilised in 0.2% (v/v) NP40 in PBS for 5 min, followed by blocking for 30 min with 1% (w/v) BSA in PBS. Permeabilised cells were stained for 1 hr with αFLAG or αPINK1 pSer228 primary antibody at indicated dilutions, followed by three washes with 0.2% (w/v) BSA in PBS. Cells were then stained for 1 hr with αMouse and αRabbit secondary antibodies conjugated to Alexa 488 and Alexa 568 respectively, at a 1:500 dilution. Following secondary antibody incubation, coverslips were washed three times in 0.2% (w/v) BSA in PBS and mounted onto slides using ProLong Gold Antifade (ThermoFisher). Images were collected on an LSM710 laser scanning confocal microscope (Cal Zeiss) using the x63 Plan-Apochromat objective (NA 1.4) and pinhole chosen to provide a uniform 0.8 μm optical section thickness in all fluorescence channels.

### Recombinant expression of phosphorylated PINK1 by genetic code expansion

Site-specific incorporation of phosphoserine into recombinant PINK1 was performed by a protocol described by Rogerson et al(17). pNHD1.3 constructs expressing MBP-HsPINK1_123-581_-6His were designed such that desired phosphoserine incorporation sites were mutated to an amber stopcodon (“*tag*”). Constructs were co-transformed into BL21 (DE3) ΔSerB cells with the pKW2 plasmid encoding the orthogonal tRNA/tRNA-synthetase pair (SepRS(2)/pSer-tRNA(B4)CUA) and the optimised elongation factor EF-Sep(17). Cells were grown in TB medium with 50% normal working concentration of antibiotic and supplemented with 2 mM O-phospho-L-serine (Sigma) at 37^ο^C and 180 rpm. At OD_600_ = 0.3, culture temperature was reduced to 18°C, and at OD_600_= 0.5, protein expression induced for 16 hrs with 250 μM IPTG.

### Recombinant PINK1 purification

16 hrs post-induction, cells were harvested and resuspended in lysis buffer (50 mM Tris-HCl pH 7.5, 150 mM NaCl, 5% (v/v) glycerol, 1 mM benzamidine, 0.1 mM PMSF, 1% Triton X-100, 0.1% (v/v) β-mercaptoethanol). Cell suspensions were further disrupted by sonication (10 rounds of 10 sec on / 10 sec off at 40% amplitude) and resulting lysates clarified by centrifugation at 30,000 xg, 4°C.

MBP-PINK1 proteins were purified from the supernatant by amylose affinity chromatography (90 min incubation with amylose resin, 500 μL per litre of culture, MRC PPU Reagents and Services). Non-specifically bound proteins were removed by four, 100x bead-volume washes in MBP wash buffer (50 mM Tris-HCl pH 7.5, 500 mM NaCl, 5% (v/v) glycerol, 1 mM benzamidine, 0.1 mM PMSF, 0.03% Brij-35, 0.1% (v/v) β-mercaptoethanol). MBP-PINK1 was eluted in 1 bead-volume of MBP elution buffer (50 mM Tris-HCl pH 7.5, 150 mM NaCl, 5% (v/v) glycerol, 1 mM benzamidine, 0.1 mM PMSF, 0.03% Brij-35, 0.1% (v/v) β-mercaptoethanol, 12 mM maltose).

6His-tagged PINK1 proteins were purified by Ni^2+^-affinity chromatography (90 min incubation with Ni^2+^-NTA resin, 500 μL per litre of culture, MRC PPU Reagents and Services). Non-specifically bound proteins were removed by four, 100x bead-volume washes in His wash buffer (50 mM Tris-HCl pH 7.5, 500mM NaCl, 5% (v/v) glycerol, 1 mM benzamidine, 0.1 mM PMSF, 0.03% Brij-35, 50 mM imidazole). 6His-tagged PINK1 was eluted in 1 bead-volume of His elution buffer (50 mM Tris-HCl pH 7.5, 500mM NaCl, 5% (v/v) glycerol, 1 mM benzamidine, 0.1 mM PMSF, 0.03% Brij-35, 200 mM imidazole). 6His-SUMO tags were cleaved by the addition of 6His-SENP1 at a ratio of 10:1 PINK1: protease, during overnight dialysis into size-exclusion buffer (50 mM Tris-HCl, 150 mM NaCl, 5% (v/v) glycerol, 1 mM benzamidine, 0.1 mM PMSF, 0.03% Brij-35).

### *In vitro* kinase assays

Reactions were set up in a final volume of 40 μl, with 500 nM of recombinant human PINK1 and 2 μM of GST-Parkin_1-76_, in 50 mM Tris–HCl pH 7.5, 0.1 mM EGTA, 10 mM MgCl_2_, 2 mM DTT and 0.1 mM [γ-^32^P] ATP (approx. 500 cpm pmol^−1^). Assays were incubated at 30°C with shaking at 1050 r.p.m. and terminated after 10 min by addition of SDS sample buffer. Reaction mixtures were then resolved by SDS–PAGE. Proteins were detected by Coomassie staining, and gels were imaged using an Epson scanner and dried completely between cellophane using a gel dryer (Bio-Rad). Incorporation of [γ-^32^P] ATP into substrates was analysed by autoradiography using Amersham hyperfilm and quantified by Cerenkov counting of respective SDS-PAGE gel bands.

### LCMS analysis of PINK1 digests using ZenoTOF 7600

PINK1 tryptic/LysC mix digests were analyzed using a Waters M Class UPLC system coupled to ZenoTOF 7600 mass spectrometer (Sciex). Digests were chromatographed on a Kinetix XB C18 150 × 0.3 mm column (Phenomenex) at 6 ml/min (A = 0.1% formic acid in water, B = 0.1% formic acid in acetonitrile) with a 21 min gradient (3–30% B) followed by 3 min (30– 80% B). The column was connected directly to the ZenoTOF 7600 OptiflowTM source fitted with a low microflow probe and heated at 30°C with the integral column oven.

LCMS data was acquired using optimal source conditions and the data dependent acquisition was performed on the Top 30 precursors (*m/z* 400–2000) with charge state 2–4+, with a minimum intensity of 300 cps and excluded for 6 sec after one occurrence. TOFMSMS spectra (*m/z* 100–2000) were acquired with Electron Activated Dissociation (EAD) using a filament current of 3000 nA, 0 eV kinetic energy, a reaction time of 10 ms and total MSMS accumulation time of 15 ms. Data files (wiff) were converted to mgf files using the AB Sciex MS Data converter (Sciex) and searched using Mascot 2.6 against Swissprot (2019 11. fasta) database. Enzyme cleavage allowed for one missed cleavage. Carbamidomethylation of cysteine was a fixed modification, phosphorylation of cysteine, serine, threonine, tyrosine, and oxidation of methionine were variable modifications. The instrument type chosen to match the EAD spectra was EThcD and the mass accuracy for precursors (20 ppm) and for MSMS spectra (0.05 Da).

## Acknowledgements

We thank Jason Chin at the MRC Laboratory of Molecular Biology for hosting A.D.W. for a Medical Research Council (MRC) funded secondment to train in gene codon expansion methods. We thank Nicole Polinski and Shalini Padmanabhan at the Michael J Fox Foundation for advice on PINK1 tool development. We are grateful to the sequencing service (School of Life Sciences, University of Dundee); James Hastie for expression and generation of recombinant proteins (MRC PPU); the MRC PPU tissue culture team (co-ordinated by Edwin Allen) and MRC PPU Reagents and Services antibody teams (co-ordinated by James Hastie). This work was supported by a Wellcome Trust Senior Research Fellowship in Clinical Science (210753/Z/18/Z to M.M.K.M.); the Michael J. Fox Foundation (M.M.K.M.), EMBO YIP Award (M.M.K.M.), Medical Research Council (M.M.K.M. and MC_UU_00028/6 to A.J.W.) and Biotechnology and Biological Sciences Research Council (M.M.K.M. and BB/R000182/1 to C.E.E.) and NorthWest Cancer Research (to C.E.E).

## Conflict of Interest

M.M.K.M. is a member of the Scientific Advisory Board of Mitokinin Inc and consultant to Stealth Biotherapeutics Inc. and MSD.

---

**Figure 1 – figure supplement 1.**
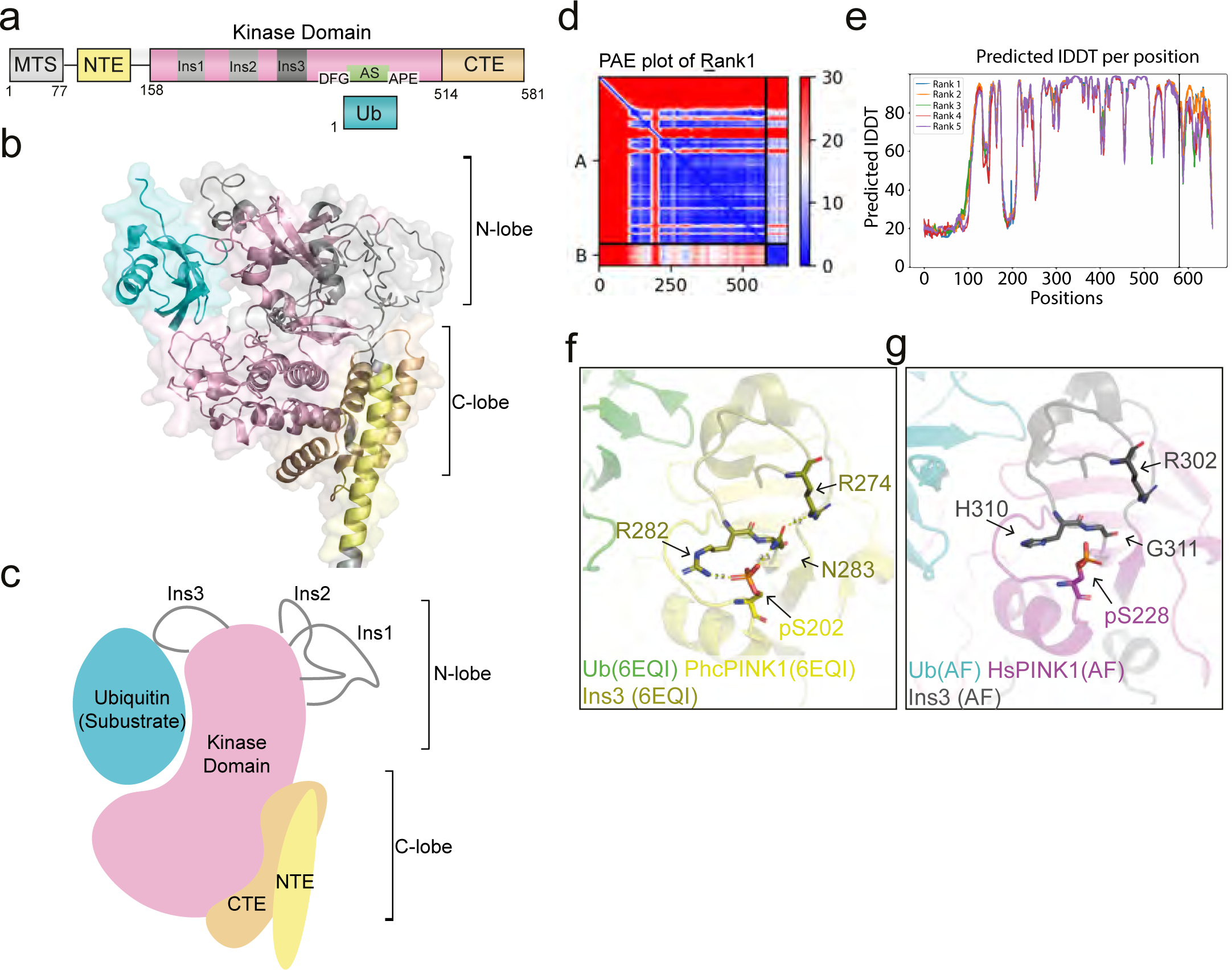
Generation of human PINK1-Ub model by AlphaFold. **(a)** Domain architecture of human PINK1. **(b)** PINK1-Ub model was generated by AlphaFold multimer v2.3.1 in Colab notebook and top-ranked model was used for further analysis. The coloring scheme applied to the PINK1-Ub model corresponds to the domain colors as illustrated in the schematic representation of the protein’s domain arrangement. **(c)** Schematic representation of domain rearrangement in PINK1-Ub model. **(d)** Predicted Aligned Error (PAE) plot indicating the confidence of each monomer; PINK1 and Ub, along with the PINK1-Ub complex packing confidence. The lines in plot create areas indicating confidence of PINK1 (top left) and Ub (bottom right) along with PINK-Ub packing dimer confidence (top right and bottom left). **(e)** Predicted local distance difference test (pLDDT) plot showing all 5 models generated by AlphaFold. The higher IDDT score indicate properly ordered protein regions. Top-ranked Model (rank 1) was for further analysis. **(f)** Detailed view of Ins3 co-ordination by autophosphorylation in the crystal structure of PhcPINK1 (PDB ID: 6EQI) [23]. **(g)** Detailed view of the equivalent region in a human PINK1 AlphaFold model.

**Figure 1 – figure supplement 2.**
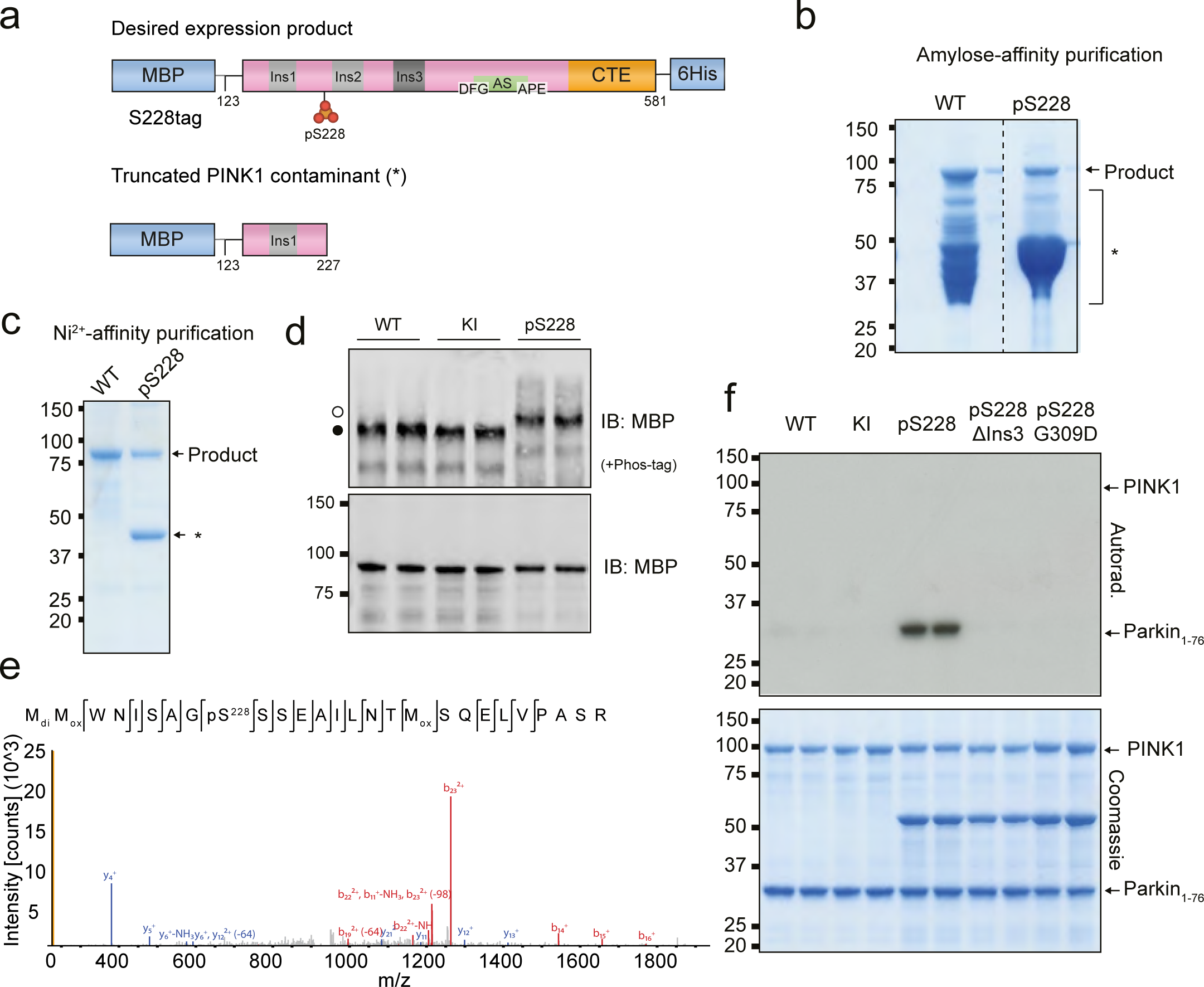
Production of recombinant human PINK1 stoichiometrically phosphorylated at Ser228 using genetic code expansion. **(a)** Construct design and expected outcomes for the desired and truncated expression product from genetic code expansion with BL21 ΔSerB +pKW2. MBP-PINK1(123-581)-6His constructs were generated, with the Ser228 codon mutated to tag for phosphoserine incorporation. Desired expression construct is a full-length sequence spanning MBP to 6His tag with pSer228 incorporated. Truncated product, resulting from early termination at the tag codon, spans only MBP tag and up to residue 227 of PINK1. **(b)** Coomassie gel of expression product from genetic code expansion (pS228) compared to the wildtype construct for no phosphorylation (WT), following an initial round of affinity purification on amylose resin. The desired expression product of ∼100 kDa is indicated, alongside the primary truncated waste productions. **(c)** Resulting proteins from amylose affinity purification in (b) were subjected to a second round of affinity purification on Ni^2+^ resin. Labelled as in (b). **(d)** Phos-tag SDS-PAGE analysis of WT and KI PINK1 from normal recombinant expression techniques compared to pSer228-loaded for SDS-PAGE, supplemented with 75 μM Phos-tag reagent, followed by αMBP immunoblot. Filled circles indicate non-phosphorylated species; unfilled circles indicate phosphorylated form of PINK1. **(e)** Confirmation of site-specific incorporation of pSer228 on the resulting recombinant protein from (c) by LC-MS/MS with CID analysis. **(f)** Full-length Coomassie-stained gel and autorad for quantification of PINK1 activity, as shown in cropped form in Figure 1d.

**Figure 1 – figure supplement 3.**
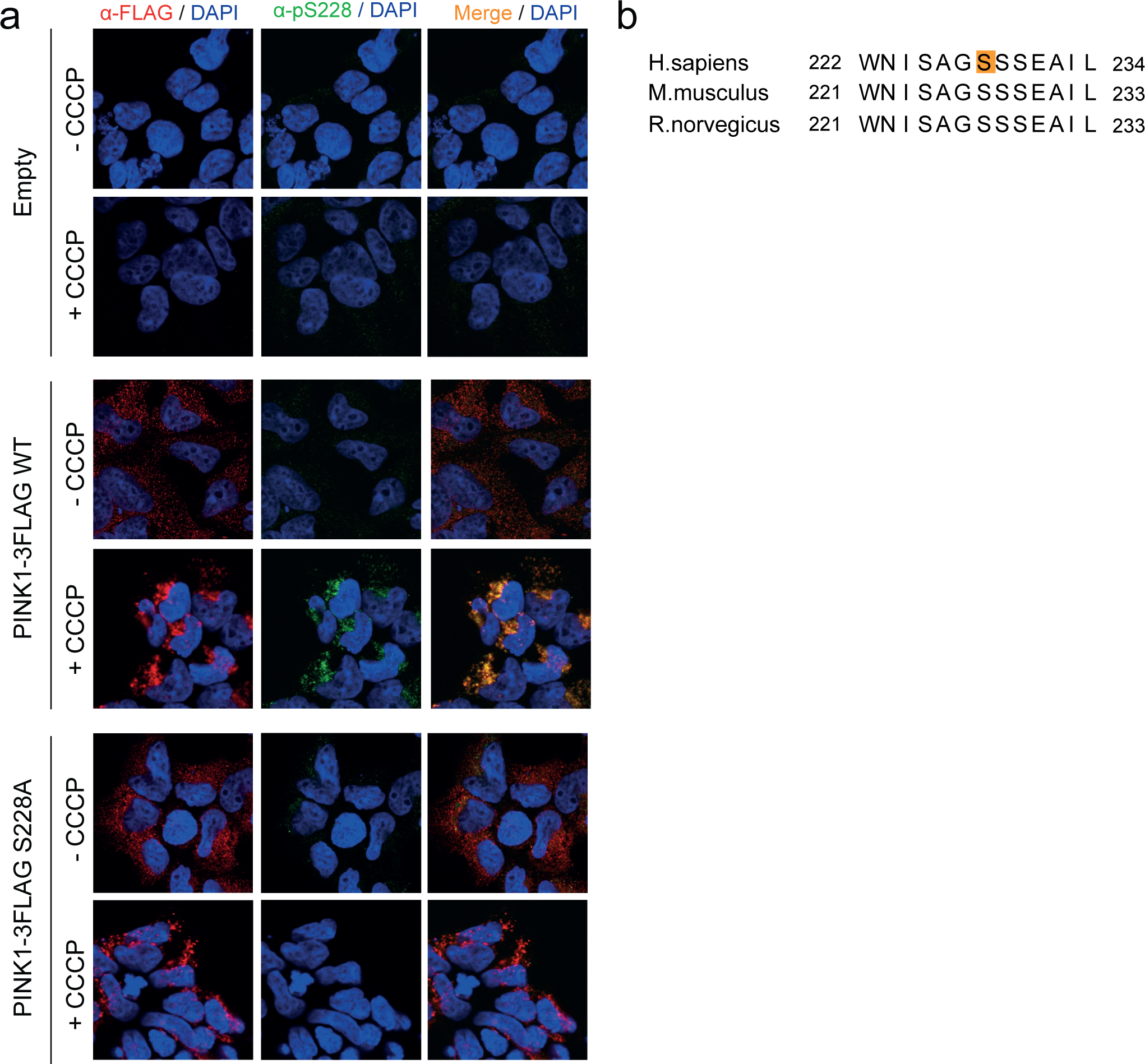
Characterisation of a pSer228-specific PINK1 antibody for immunofluorescence. **(a)** Testing of antibody against native protein by immunofluorescence. PINK-knockout HEK293 cells stably re-expressing PINK1-3FLAG WT or S228A were seeded on coverslips and treated with doxycycline and CCCP as in (a). Cells were fixed with 4% (v/v) paraformaldehyde and probed with αFLAG and αpSer228 antibodies at 1 μgmL^-1^, then visualised by Alexafluor-488 and 568-coupled secondary antibodies, respectively. (**b)** Sequence alignment of the pSer228 epitope from human PINK1 relative to mouse and rat PINK1. Alignments were performed in MUSCLE and annotated in JalView.

**Figure 2 – figure supplement 1.**
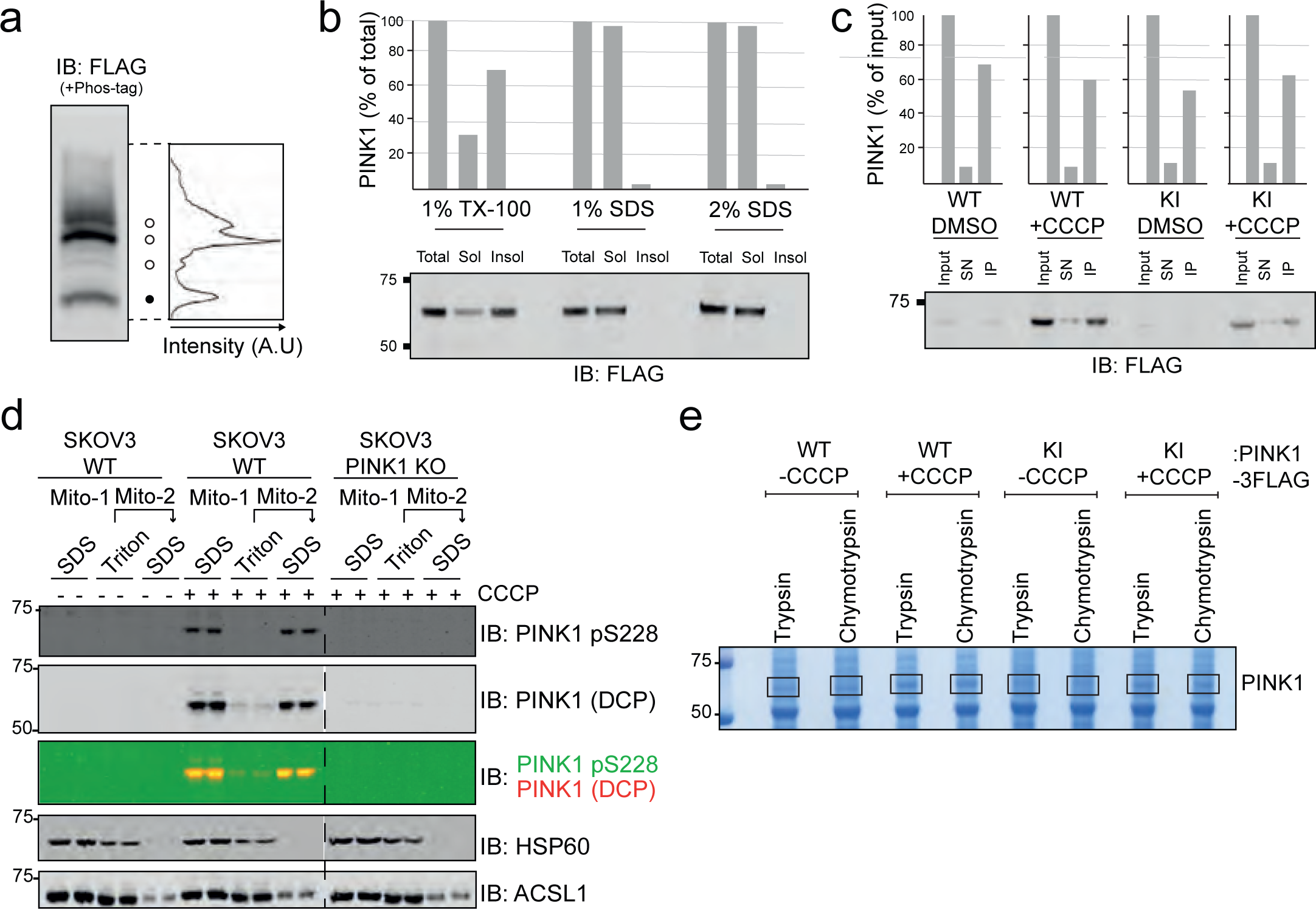
Sample preparation for phosphoproteomics. **(a)** Detailed view of Phos-tag immunoblot (IB) for PINK1-3FLAG WT. Filled circles indicated non-phosphorylated species; unfilled circles indicate the two distinct phosphorylated species. Licor densitometry for the 3 total PINK1 species is shown. **(b)** Optimisation of PINK1-3FLAG WT extraction from mitochondrial fractions using lysis buffer supplemented with indicated detergents. For each detergent, the post-lysis soluble (sol) and insoluble (insol) fractions are quantified relative to a pre-lysis (total; 100%) sample. **(c)** Quality control for immunoprecipitation efficiency for PINK1-3FLAG. 10 mg of mitochondrial extracts from indicated HEK293 cell lines (expressing PINK1-3FLAG WT or KI, treated or untreated with CCCP) were lysed in 1% SDS and diluted down to a final SDS concentration of 0.1%, then PINK1 immunoprecipitated on αFLAG-agarose. Resulting immunoprecipitates were diluted such that total PINK1-3FLAG content could be directly compared to the pre-IP (input) and unbound supernatant (SN) fractions. **(d**) The presence of endogenous PINK1 in the mitochondrial fraction of WT and PINK-1 KO SK-OV-3 cells was assessed using Triton or SDS lysis. A mitochondrial pellet of 100 µg for each condition was either lysed directly in 1% SDS or initially in 1% Triton followed by lysis of the remaining pellet in 1% SDS. The samples were then examined for the presence of pSer228 and PINK1, which were predominantly detected in the SDS-solubilized pellet, in contrast to the matrix protein HSP-60 or the outer-mitochondrial membrane protein ACSL1. **(e)** SDS-PAGE analysis of immunoprecipitates. Immunoprecipitated samples from 10 mg mitochondria were run in duplicate on 4-12% SDS-PAGE gels, for each condition as in (c). Gels were stained with Coomassie Blue and bands corresponding to PINK1 (boxed) were excised for digestion with either trypsin or chymotrypsin.

**Figure 2 – figure supplement 2.**
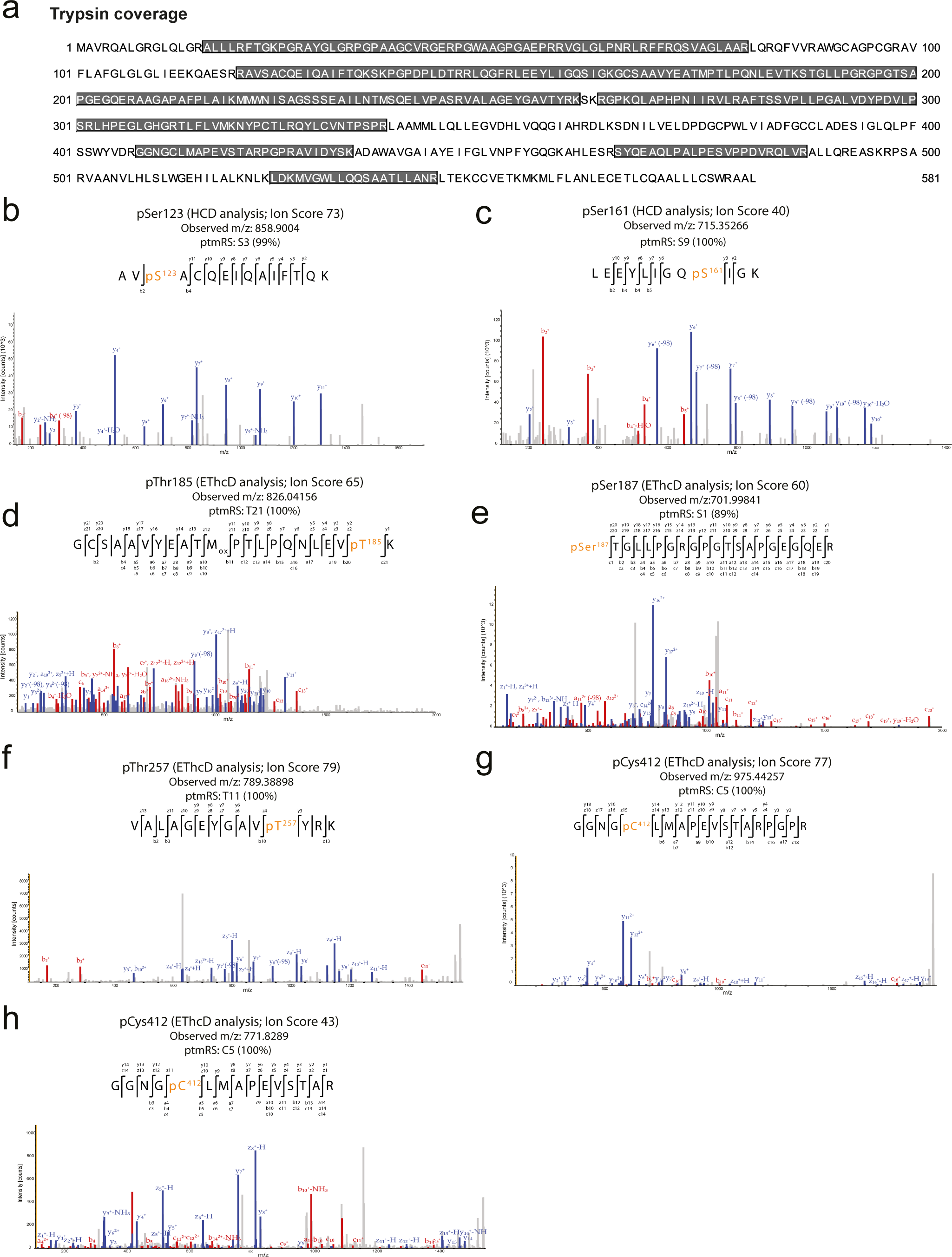
Identification of PINK1-3FLAG Ser/Thr and Cys phosphorylation sites of hPINK1 from HEK293 cells following digestion with trypsin. **(a)** Sequence coverage of tryptic peptides. Found peptides are highlighted in grey. **(b-h)** MS2 spectrum, observed *m/z* and ptmRS score **(b)** Tandem mass spectrum of the doubly charged ion at *m/z* 858.9004 following HCD fragmentation with a Mascot ion score of 73 indicates phosphorylation at Ser123. Tryptic peptide containing the Ser123 phosphorylation site (ptmRS: 99%), following fragmentation by higher-energy collision dissociation (HCD). **(c)** Tandem mass spectrum of the doubly charged ion at *m/z* 715.35266 following HCD fragmentation with a Mascot ion score of 40 indicates phosphorylation at Ser161. Tryptic peptide containing the Ser161 phosphorylation site (ptmRS: 100%), following fragmentation by HCD. −98 represents the loss of H_3_PO_4_. **(d)** Tandem mass spectrum of the doubly charged ion at *m/z* 826.04156 following EThcD fragmentation with a Mascot ion score of 65 indicates phosphorylation at Thr185. Tryptic peptide containing the Thr185 phosphorylation site (ptmRS: 100%), following fragmentation by Electron-transfer/higher-energy collision dissociation (EThcD). **(e)** Tandem mass spectrum of the triply charged ion at *m/z* 701.99841 following EThcD fragmentation with a Mascot ion score of 60 indicates phosphorylation at Ser187. Tryptic peptide containing the Ser187 phosphorylation site (ptmRS: 89%), following fragmentation by EThcD. **(f)** Tandem mass spectrum of the doubly charged ion at *m/z* 789.38898 following EThcD fragmentation with a Mascot ion score of 79 indicates phosphorylation at Thr257.Tryptic peptide containing the Thr257 phosphorylation site (ptmRS: 100%), following fragmentation by EThcD. **(g)** Tandem mass spectrum of the doubly charged ion at *m/z* 975.44257 following EThcD fragmentation with a Mascot ion score of 77 indicates phosphorylation at Cys412. Miscleaved tryptic peptide containing the Cys412 phosphorylation site (ptmRS: 100%), following fragmentation by EThcD. **(h)** Tandem mass spectrum of the doubly charged ion at *m/z* 771.82892 following EThcD fragmentation with a Mascot ion score of 43 indicates phosphorylation at Cys412. Tryptic peptide containing the Cys412 phosphorylation site (ptmRS: 100%), following fragmentation by EThcD.

**Figure 2 – figure supplement 3.**
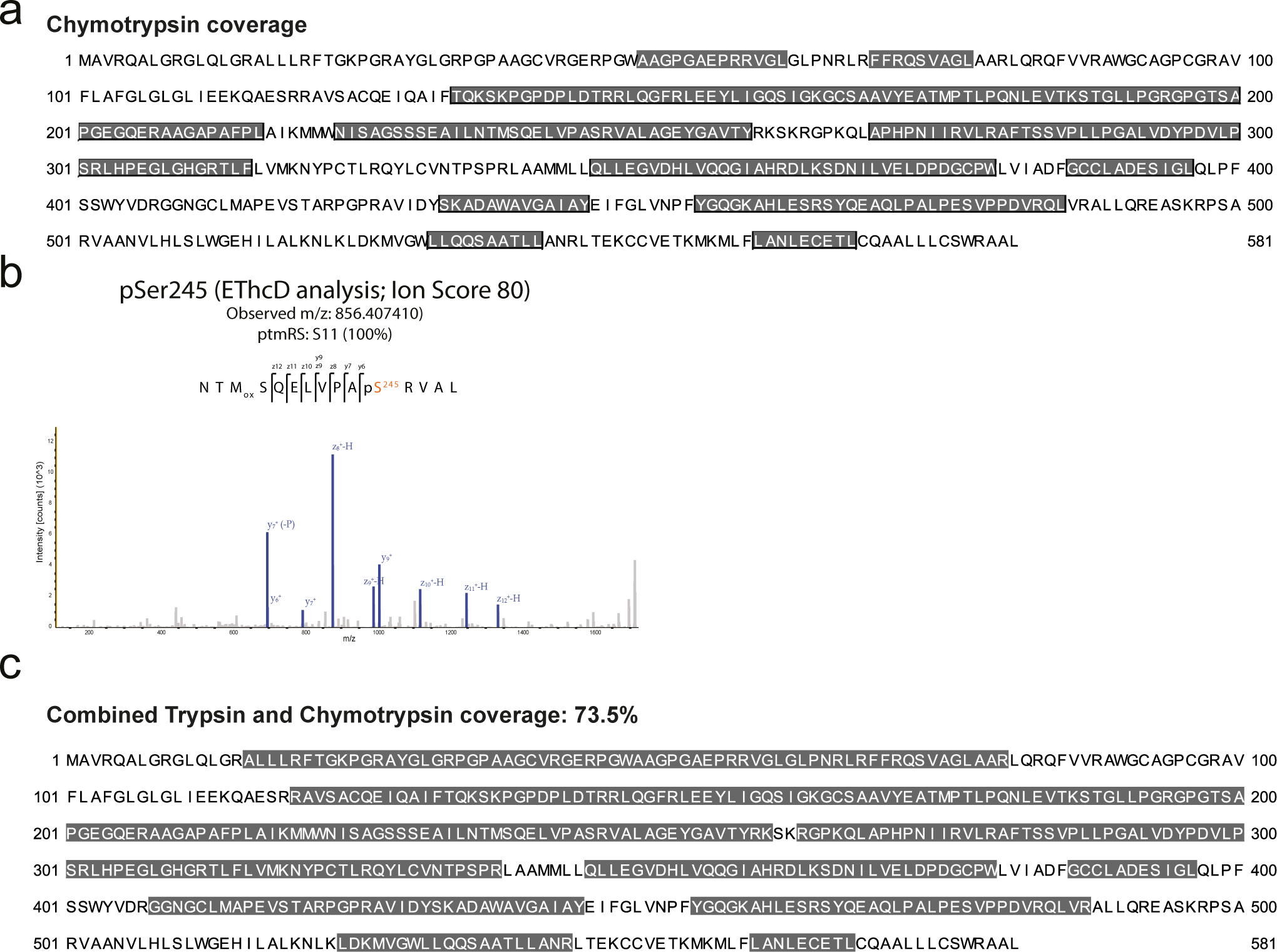
Identification of PINK1-3FLAG Ser/Thr autophosphorylation sites of hPINK1 from HEK293 cells following digestion with chymotrypsin. **(a)** Sequence coverage of chymotryptic peptides. Annotated in JalView. **(b)** Tandem mass spectrum of the doubly charged ion at *m/z* 856.40741 following EThcD fragmentation with a Mascot ion score of 80 indicates phosphorylation at Ser245. MS2 spectrum, observed *m/z* and ptmRS score of 100% corresponding to chymotryptic peptide containing the Ser245 phosphorylation site, following fragmentation by electron-transfer/higher-energy collision dissociation (EThcD). −P represents the loss of H_3_PO_4_. **(c)** Sequence coverage of combined trypsin and chymotryptic peptides. Annotated in JalView.

**Figure 2 – figure supplement 4.**
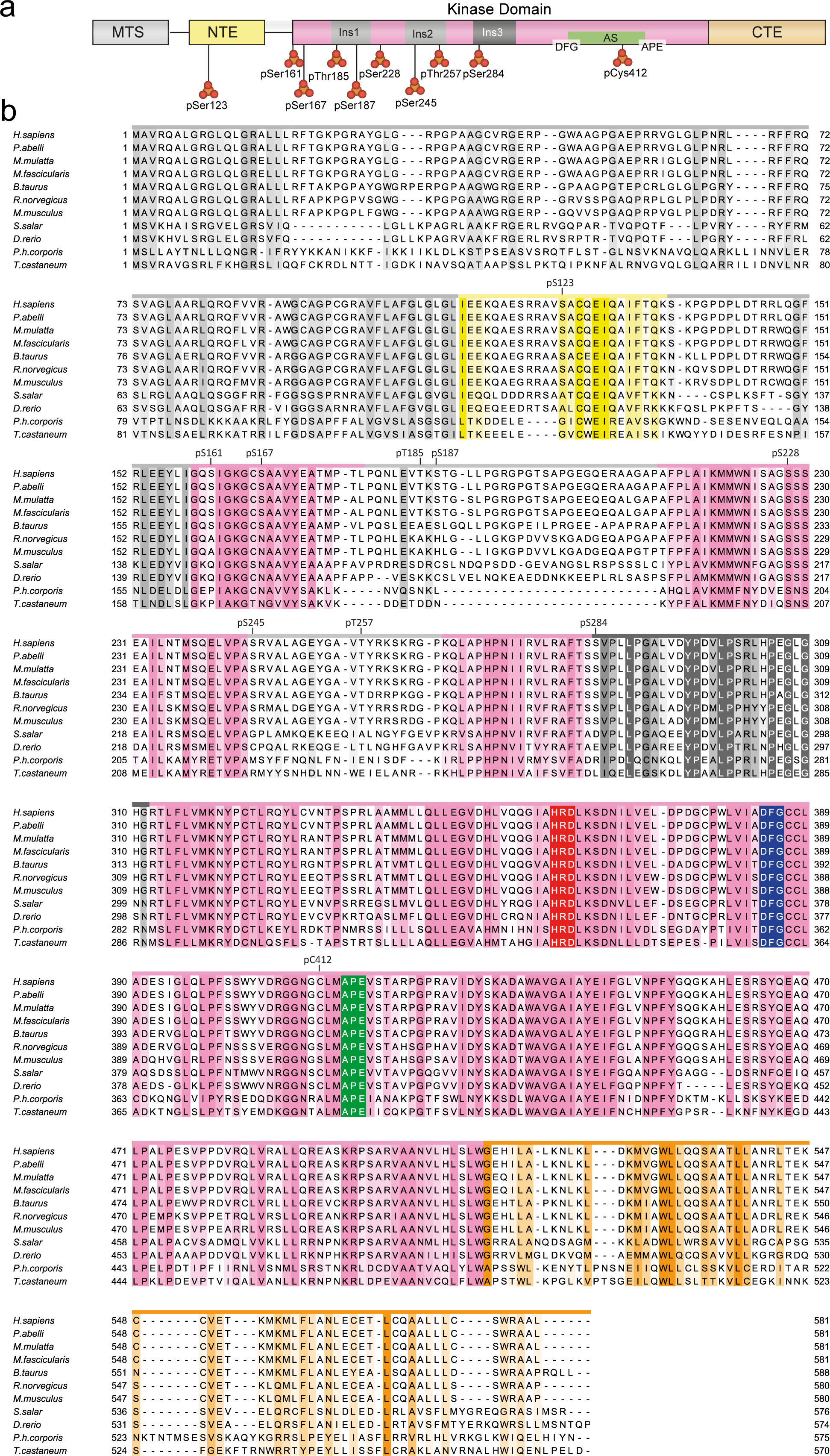
Multiple sequence alignment of PINK1 species. **(a)** Domain architecture of hPINK1 with mitochondrial targeting sequence (MTS), N-terminal extension (NTE), activation segment (AS), C-terminal extension (CTE), and all identified autophosphorylation sites indicated. **(b)** Multiple sequence alignment of PINK1 primary sequence across select eukaryotic species. Alignments were performed in MUSCLE and annotated in JalView. Coloured according to conservation and domain as illustrated in the schematic representation of the protein’s domain arrangement.

**Figure 2 – figure supplement 5.**
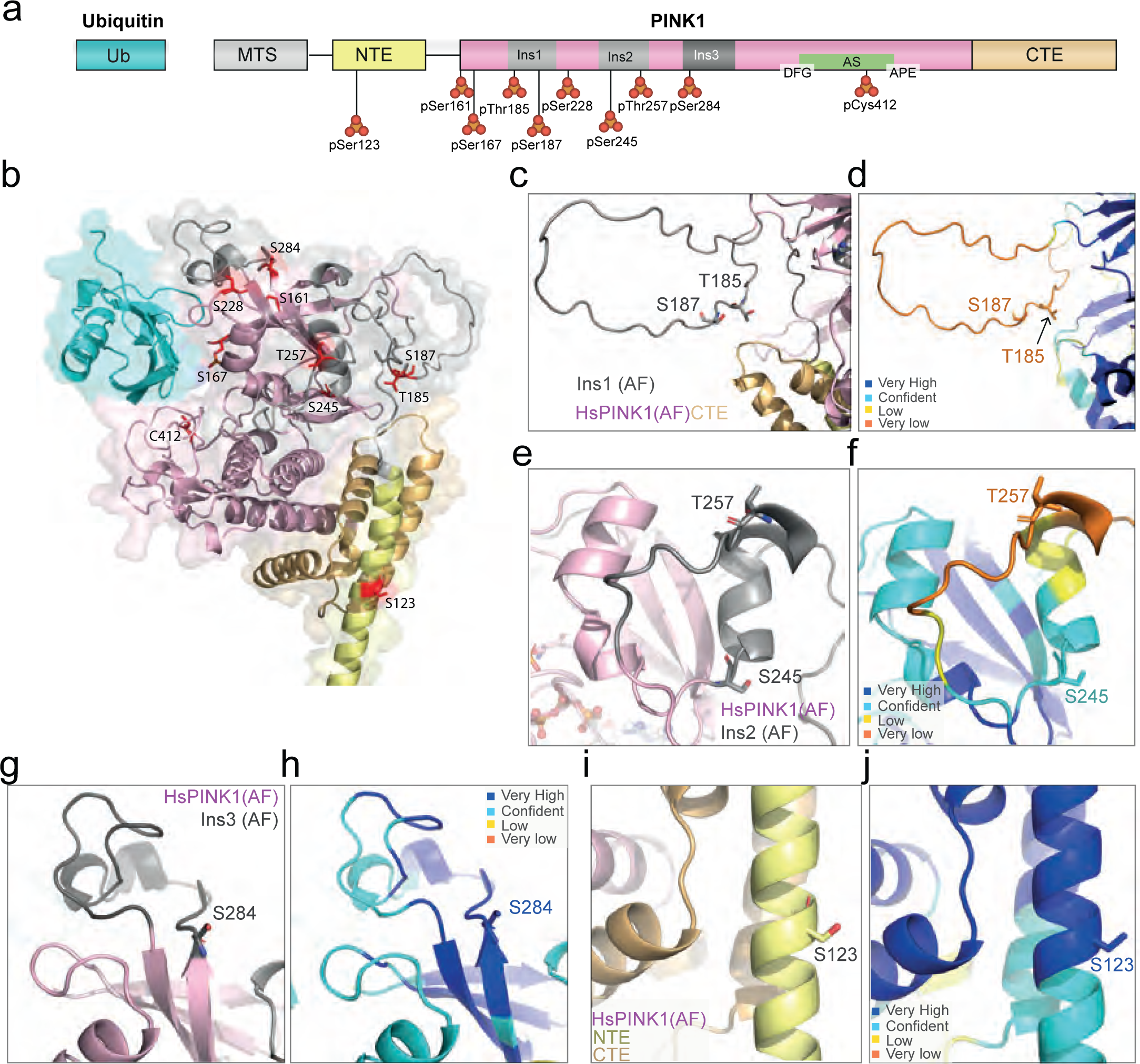
Phosphorylation sites on human PINK1 model. **(a)** Domain architecture of hPINK1 with all identified autophosphorylation sites indicated. **(b)** All identified autophosphorylation sites on Human PINK1-Ub AlphaFold model. **(c,e,g,i)** Detailed view of autophosphorylation sites in PINK1 AlphaFold model. The coloring scheme applied to the PINK1-Ub model corresponds to the domain colors as illustrated in the schematic representation of the protein’s domain arrangement. **(d,f,h,j)** Prediction confidence of residues in (c,e,g,i), respectively, on the AlphaFold model of PINK1. Regions are coloured according to the pLDDT score.

**Figure 4 – figure supplement 1.**
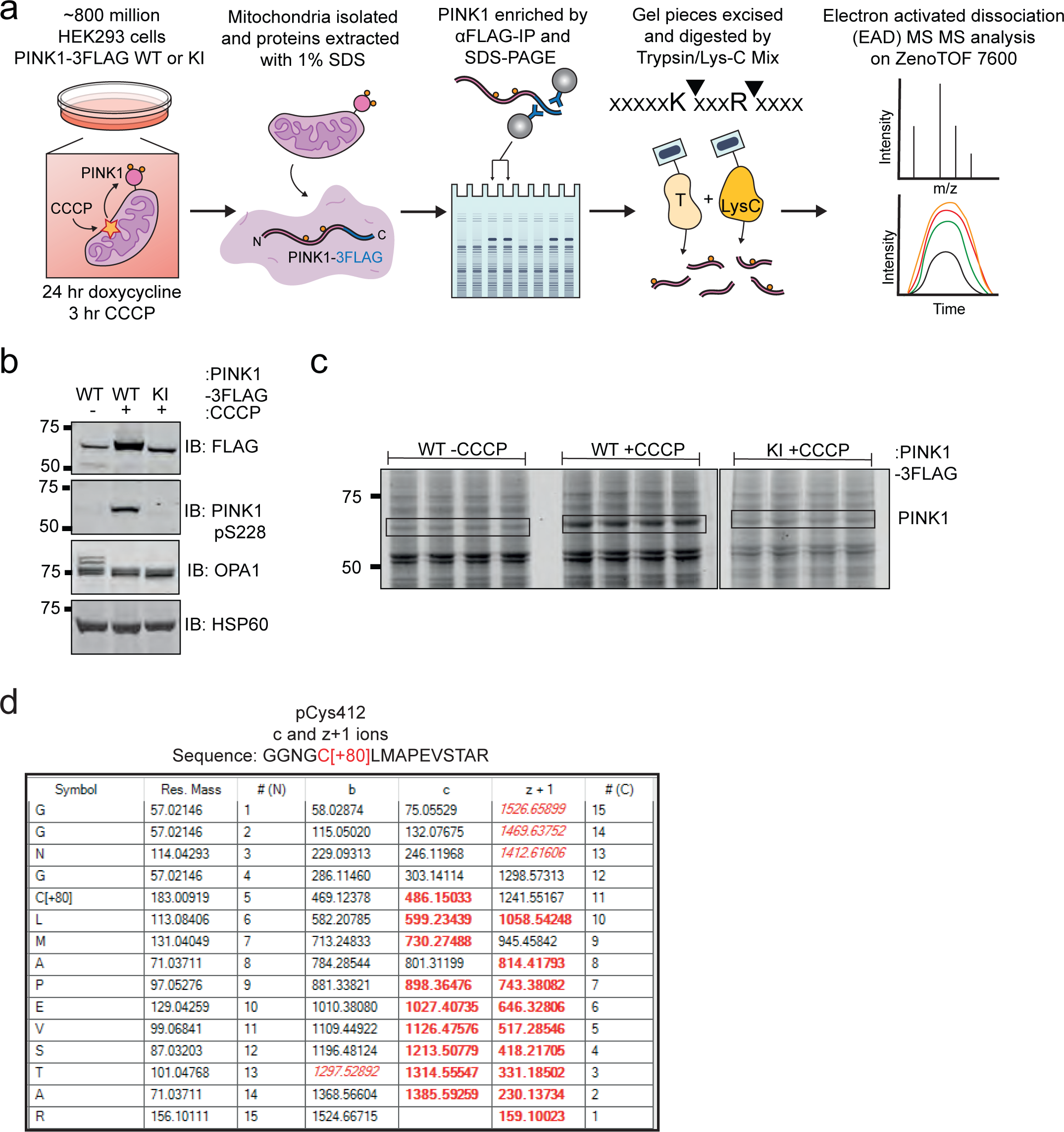
Workflow for phosphoproteomic analysis on ZenoTOF 7600. **(a)** Workflow of sample preparation for human PINK1 EAD MS/MS analysis. ∼800 million HEK293 cells were treated with doxycycline for 24 hr followed by treatment with CCCP in the last 3 hr. Mitochondrial fractions were isolated and lysed in 1% SDS buffer and PINK1 was immunoprecipitated with αFLAG-amylose resin, then resolved on 4-12% gradient acrylamide gels by SDS-PAGE. Gel pieces were excised and subjected to in-gel digestion with Trypsin+LysC mix for EAD MS/MS analysis on ZenoTOF 7600. **(b)** Quality control blot for the mitochondrial extracts from PINK1-3FLAG WT or, KI which were subjected to PINK1 immunoprecipitation with αFLAG-amylose resin. **(c)** SDS-PAGE analysis of immunoprecipitates. Immunoprecipitated samples from 20 mg mitochondrial extracts were run on 4-12% SDS-PAGE gels divided in 4 lanes for each condition and gels were stained with Coomassie Blue and bands corresponding to PINK1 (boxed) were excised for digestion with Trypsin+LysC mix. **(d)** EAD MSMS spectrum table Cys412. All matched c and z+1 ions of GGNGpCLMAPEVSTAR (*m/z* 514.9 3+) are highlighted in red in the table. Phosphocysteine is annotated as C[+80].

**Figure 4 – figure supplement 2.**
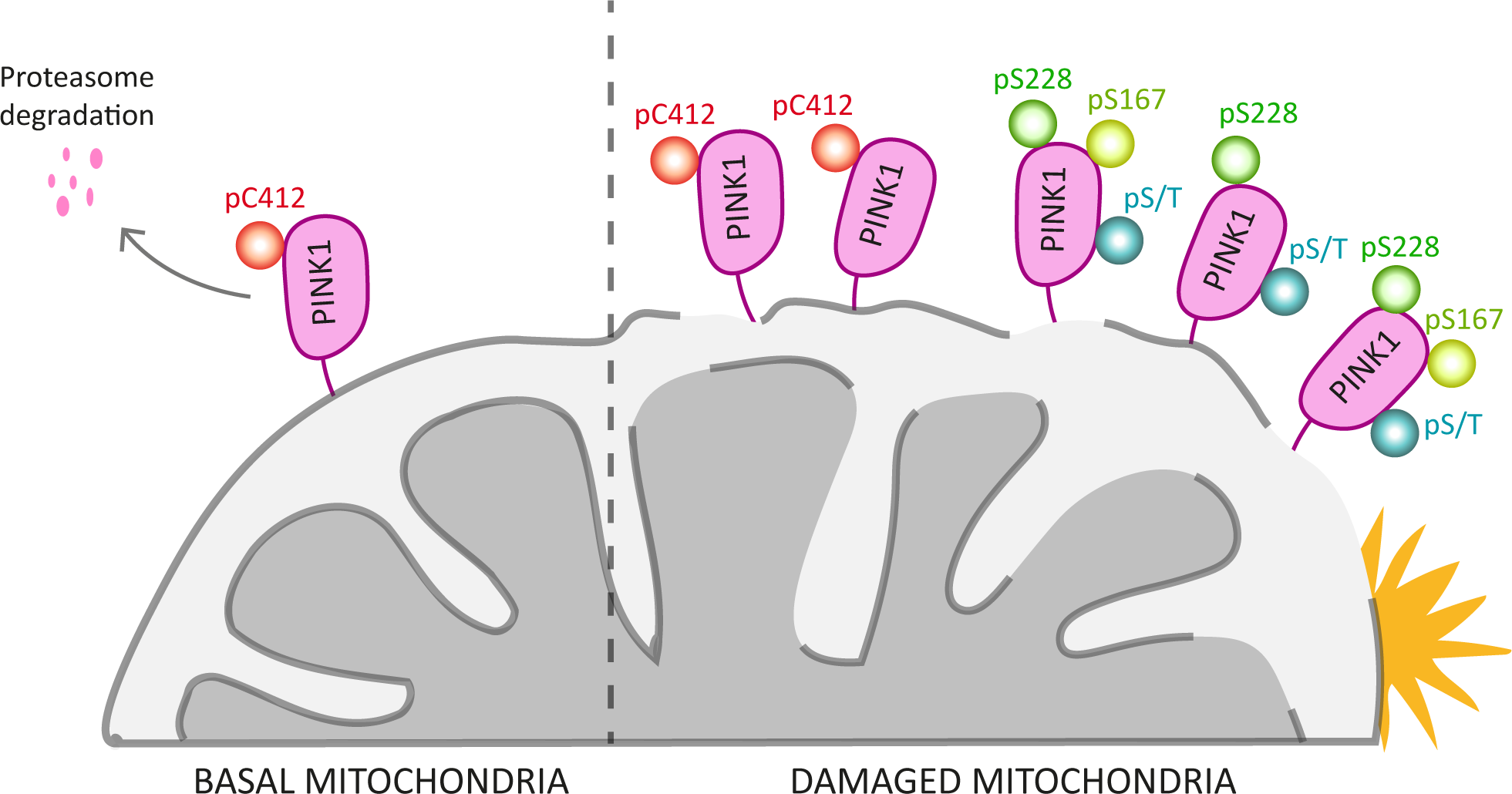
Model of phospho-*h*PINK1. Model of phosphorylation status of *h*PINK1 under basal and mitochondria damage. Our findings suggest that phosphorylation of Cys412 functions mainly as a negative regulator of *h*PINK1. Upon mitochondrial damage, *h*PINK1 accumulates due to stabilisation of pre-formed *h*PINK1 and new translation of *h*PINK1 that undergoes auto-phosphorylation at multiple Ser/Thr sites. The activatory Ser228 and Ser167 sites are indicated.

